# Network reconfiguration preserves prediction error signalling in the aging brain

**DOI:** 10.64898/2026.04.11.717798

**Authors:** Mathias Houe Andersen, Gemma Fernández-Rubio, David R. Quiroga-Martinez, Mattia Rosso, Mathias Klarlund, Kit Melissa Larsen, Hartwig Roman Siebner, Morten L. Kringelbach, Peter Vuust, Leonardo Bonetti

## Abstract

Cognitive aging is widely associated with a progressive weakening of predictive brain mechanisms. This view is supported by decades of electrophysiological studies reporting attenuated mismatch responses in older adults. Yet the literature remains inconsistent, suggesting that aging may not uniformly attenuate predictive processing. One possibility is that multiple predictive subsystems operate concurrently but have rarely been disentangled. Here we address this question by separating whole-brain networks underlying predictive processing in source-reconstructed magnetoencephalography (MEG) data from 77 younger and older adults performing the auditory local–global paradigm. Network decomposition revealed three temporally overlapping predictive subsystems with distinct functional profiles. Aging exerted selective effects across these networks. Sensory prediction error responses were enhanced within a network linking auditory cortices with medial cingulate regions, whereas responses associated with reorientation of attention and contextual pattern processing were attenuated in older adults. The level of multivariate recurrency across these networks was preserved with aging, while the processing of sensory violations induced more recurrency and less divergence relative to contextual violations in both groups. These findings challenge the prevailing view that predictive processing simply declines with age. Instead, aging redistributes predictive resources across distinct neural systems, amplifying sensory-based processes while weakening more cognitively demanding predictive mechanisms.

## Introduction

Predictive coding posits that perception arises from the brain’s continuous attempts to anticipate incoming sensory signals and that internal models are updated when sensory evidence violates those expectations. Such mismatches generate prediction errors at multiple hierarchical and temporal scales within the nervous system^1–6^. In the auditory domain, predictive processing spans a broad range, from local predictions about immediate tone features to global expectations about abstract, higher-order regularities unfolding over extended time windows^7–9^.

Age-related changes in predictive processing have been studied most extensively at the local sensory level. The prevailing view is that aging weakens prediction error signalling, particularly for simple sensory violations. This interpretation is supported by numerous electrophysiological studies reporting reduced mismatch responses in older adults and has often been framed as an age-related shift in the balance between top-down priors derived from internal models and bottom-up evidence sampled through sensory input channels^10–18^. As sensory systems deteriorate with age, incoming signals may become less precise, encouraging the brain to rely more heavily on prior expectations and less on noisy incoming sensory input^19,20^. In such a framework, attenuated prediction error signalling would be an adaptive consequence of reduced sensory reliability^21^.

Yet this account remains incomplete. Findings across the aging literature are not fully consistent with this account, and the evidence for a uniform decline in prediction error signalling is weaker than often assumed. Although many studies report reduced mismatch negativity (MMN) responses in older adults^17,22–25^, meta-analytic work also finds that a substantial number of experiments detect no age-related MMN reduction^16^. In several cases, preserved mismatch responses have been attributed to altered recruitment of attention-related neural processes^15,26–32^. Similarly, in the relatively few studies using paradigms that probe both local sensory violations and higher-order pattern violations, age effects have not been uniform across levels. While MMN may be preserved, later responses linked to conscious contextual updating and attentional control, such as the global prediction error (GPE) and reorientation of attention (RON) responses, appear more consistently attenuated with age^7,8,21,33–35^. This pattern raises the possibility that aging does not uniformly reduce predictive processing but rather alters how distinct predictive subsystems are engaged.

A major limitation of prior work is that most studies have relied on scalp-level magneto-or electroencephalography (M/EEG) analyses^36,37^ and on a component-centric logic focused on canonical event-related responses such as MMN, P3a, the global prediction error response, and RON^41,38–43^. These responses are useful markers, but they do not directly reveal the large-scale brain networks that generate them, nor how such networks may differ across age. Source-localisation studies suggest that local violations recruit bilateral fronto-temporo-parietal regions, including Heschl’s gyrus and surrounding auditory cortex^7,44^, whereas global violations engage broader fronto-parieto-temporal systems overlapping with global workspace accounts of conscious access^7,45–47^. However, these network-level systems have only rarely been disentangled in relation to specific predictive responses and have not been compared across age in order to characterise their temporal dynamics.

To address this gap, we apply a recently developed method, BROADband Network Estimation via Source Separation (BROAD-NESS)^48,49^, to separate simultaneous, orthogonal brain networks from source-reconstructed MEG data. This approach makes it possible to move beyond single sensor-level components and identify concurrent large-scale networks underlying predictive processing. Importantly, BROAD-NESS also enables a dynamical characterisation of these networks. By embedding the dominant component time series into a low-dimensional phase space, brain activity can be described as trajectories through a shared network state space. Recurrence quantification analysis applied to these trajectories provides quantitative indices of stability, predictability, persistence, and complexity, revealing how often the system revisits similar network states over time. This allows us to ask not only which networks support prediction error signalling, but also whether the dynamical regime of predictive processing differs between sensory and pattern violations and whether these distinctions are preserved in aging.

In the present study, we applied BROAD-NESS to source-reconstructed MEG data from younger and older adults performing the auditory local–global paradigm. Our aims were threefold: (i) to identify the dominant whole-brain networks supporting hierarchical prediction error signalling; (ii) to reconstruct group-level network time series in order to relate these networks to canonical event-related field (ERF) responses; and (iii) to quantify how the phase-space dynamics and recurrence structure of these networks differ between sensory and pattern violations and across age. By doing so, we test the conventional decline account, which posits that predictive processes attenuate with age independent of task demands and characterize the network configurations underlying these processes.

## Results

### Overview of the experimental design and MEG source reconstruction

Seventy-seven participants engaged in an adapted version of the auditory local-global paradigm^7^ during magnetoencephalography (MEG) recordings (Fig. 1a-b). The task consisted of 160 trials per block for a total of 4 blocks. Each block included a combination of patterns consisting of either five repetitive tones, or patterns where the fifth tone deviated in frequency (Fig. S1). The local-global paradigm was designed to elicit local neural responses from sensory violations at the single tone level and global neural responses from violations at pattern level across blocks. The paradigm shifted between two block types, where the number of occurrences where the fifth tone differed in frequency was changed to create a global violation of the expected pattern. In both block types, a target tone requiring a button press was also presented to ensure continued attention towards the paradigm.

**Fig. 1.**
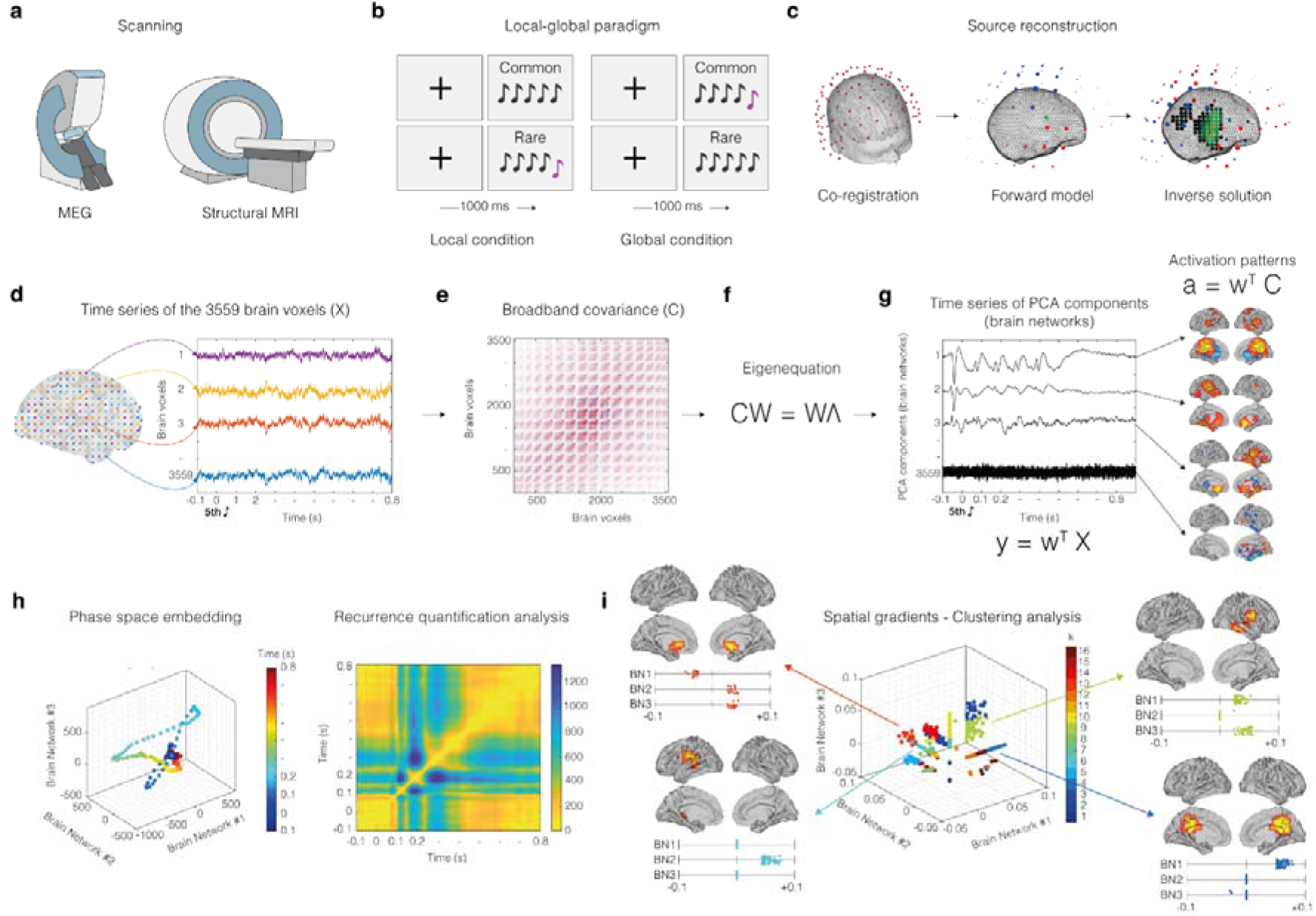
| Methodology – BROADband network estimation via source separation (BROAD-NESS). **a**, MEG was used to collect neurophysiological data while participants listened to the auditory local-global paradigm. **b**, Auditory sequences of five tones, containing either five repetitive, or four repetitive tones and a deviating fifth tone, were presented in random order. Global deviations were achieved by switching the probability of occurrences between the two sequences, producing a pattern violation. **c**, Co-registration was performed between MEG data and anatomical MRI scans of each individual. To reconstruct the neural sources that generated the signal recorded by the MEG, a single shell forward model was used. The inverse solution was estimated through beamforming. **d**, Source reconstruction yielded time series data for 3,559 brain voxels from an 8-mm grid brain parcellation. **e**, The covariance matrix (C) was computed from the broadband voxel data matrix. **f**, Principal Component Analysis (PCA) was computed by solving the eigenequation CW = WΛ for the eigenvectors (W) to find the weighted combinations of brain voxels to orthogonal components, which reflected brain networks; associated eigenvalues (Λ) express the amount of variance explained by each network component. **g**, Network activation time series (y) were derived by applying the spatial filters (w) to the voxel data matrix (X). The same filters were applied to the covariance matrix (C) to compute the spatial activation patterns (a), representing the projection of components in voxel space. **h**, Temporal embedding and recurrence quantification analysis were applied to the network time series to investigate their dynamic temporal properties. Phase space plots (left) and recurrence plots (right) captured differences in trajectory recurrency across experimental conditions. **i**, To examine the spatial organisation of voxel contributions to each brain network, we performed a spatial gradient embedding and clustering analysis based on the spatial activation patterns of the three main brain networks. An unsupervised clustering procedure applied to this space revealed anatomically interpretable voxel groups, including clusters of voxels selectively involved in one or two networks.

After performing pre-processing of the MEG data (see Methods for details), we computed source reconstruction and obtained the difference wave by subtracting the processing of standard stimuli from that of deviant stimuli before. We employed a single-shell forward model alongside a beamforming approach as the inverse solution, using an 8-mm grid that corresponded to 3,559 brain voxels (Fig. 1c-d). This procedure produced a time series for each reconstructed brain voxel returning a matrix of 3,559 brain voxels x time-points, independently for each participant and experimental condition.

### Deriving brain networks via PCA

PCA was conducted on the reconstructed time series of the 3,559 brain voxels, averaged across participants and conditions, to identify simultaneously operating broadband brain networks (Fig. 1e-f-g). Effective dimensionality (ED)^50^ was computed from the PCA eigenvalue spectrum to quantify the number of components that meaningfully contribute to the data (see Methods for details). This procedure identified three significant PCs (ED = 2.835), which accounted for 51.78 %, 24.57 % and 15.12 % of the variance in the data. Each component was interpreted as a distinct brain network based on its spatial activation pattern, which was obtained by multiplying the eigenvector of the component with the covariance matrix of the data.

As illustrated in Fig. 2, the first identified brain network comprised bilateral auditory cortices such as Heschl’s gyri, Rolandic Operculum, the Medial and Posterior Cingulate Cortices, the Precuneus and the Insula. The second network involved auditory cortex including Heschl’s gyrus and Fusiform gyrus, Hippocampus and Rolandic Operculum of the left hemisphere and bilateral frontal areas such as Rectus, Olfactory Bulb and Subgenual Anterior Cingulate Cortices (ACC). The third network showed a closely mirrored pattern of the second network, displaying connections between Heschl’s gyrus, Fusiform gyrus, Hippocampus, Rolandic Operculum, Caudate and Supramarginal gyrus of the right hemisphere and bilateral frontal areas including Rectus, Olfactory Bulb and Subgenual ACC. For specific MNI coordinates and voxel counts within each brain area, see Table S1 and S2. The remaining networks were deemed less relevant as they explained considerably less variance and fell below the effective dimensionality (ED) threshold.

**Fig. 2.**
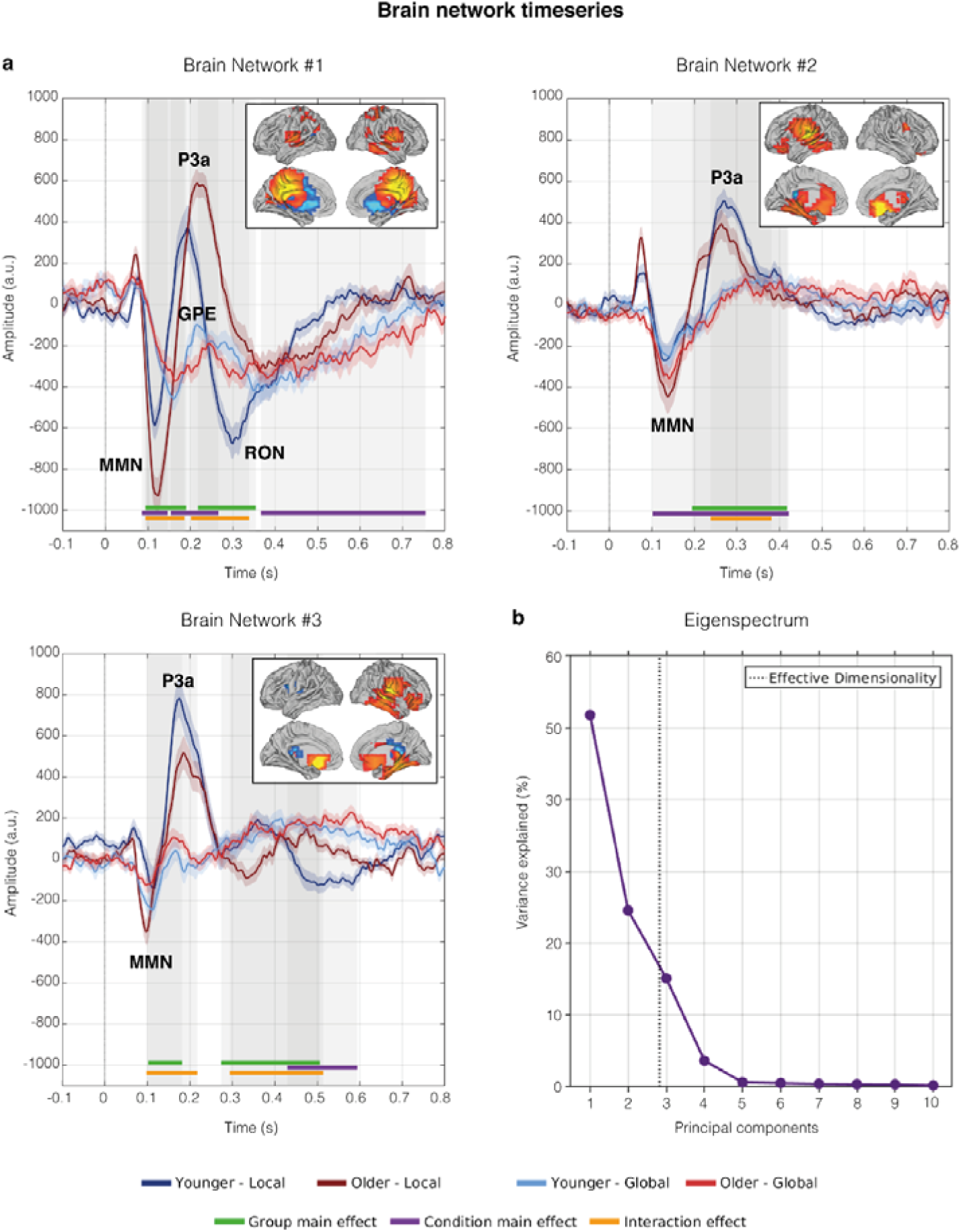
| Time series and spatial activation patterns of the main brain networks. The figure illustrates the time series and spatial activation patterns of the three brain networks that explained the highest variance (51.78 %, 24.57 % and 15.12 %, respectively). These networks were estimated using PCA within the BROAD-NESS framework, computed on the data averaged across conditions and participants. **a**, Independent time series for each participant, brain network, and experimental condition were generated using PCA-derived weights from the averaged data. The individual time series were then averaged across participants, as shown in the plots. Shaded areas represent standard errors of the mean. The brain templates illustrate the spatial extent of the networks, with yellow voxels contributing the most and light blue voxels contributing the least to the time series. Only voxels with values exceeding the mean by more than one standard deviation in absolute terms are depicted. Significant group, condition and interaction effects are indicated by green, purple and yellow lines. Time = 0 s. indicates the onset of the fifth tone in the local-global paradigm. **b**, The eigenspectrum for the first 10 principal components and effective dimensionality (ED = 2.835), indicates the relevance of the first three principal components (dashed line).

After deriving the time series in each network at participant level, statistical analyses examining age group, condition and interaction effects were conducted for each of the first three brain networks. This analysis revealed several time windows of significant differences between the local and global condition, between age groups and between the interactions of condition and age group (Fig. 2). Regarding age group effects, a general pattern was observed. In Brain Network #1, older adults displayed enhanced local MMN and P3a responses, and an attenuated RON response compared to younger adults. Furthermore, the global prediction error response was reduced in older adults. In Brain Network #2 and #3, younger adults displayed an increased P3a response, while the RON response was also increased among younger adults in Brain Network #3. Several significant condition and interaction differences were found throughout the time window. Detailed cluster onsets/offsets and cluster-level p-values are summarised in Table S3. See Fig. S2 for details regarding the detection of neural responses.

Furthermore, by examining the latency of key ERF responses across these three main brain networks in each group for each condition, a specific order of elicited responses was indicated in both the local and the global condition (Fig. S2, Table S4). In the local condition, ERF responses were firstly elicited in Brain Network #3 followed by Brain Network #1 and Brain Network #2. This indicated that local ERF responses were predominantly initiated in the right auditory cortex, followed by activation of the central network covering the cingulate cortices and finally terminating in auditory regions of the left auditory cortex. In the global condition, ERF responses were similarly initiated in Brain Network #3 covering the right auditory cortex but were then followed by Brain Network #2 covering mainly the left auditory cortex and terminating in Brain Network #1, the central network covering cingulate cortices.

### Phase space and recurrence quantification analysis

To further characterise the temporal organisation of BROAD-NESS-derived brain network dynamics, we embedded the three dominant networks into a joint phase space and quantified their recurrence structure. Visual inspection of the three-dimensional trajectories (Fig. 3a) revealed more sharply defined, time-locked utilisation of brain networks in the local condition relative to the global condition. In the local deviation condition in the time range of 150 to 400 ms post onset of the fifth tone, younger adults exhibited a stronger co-activation between all three brain networks, whereas older adults relied more heavily on Brain Network #1. When specifically examining age group differences in the three-dimensional phase space trajectories of the local condition, it became apparent, that during the timing of the early local MMN and P3a responses, older adults utilised Brain Network #1 to a larger extent, while this network was more involved for younger adults during the late local RON response (Fig. 3b). Furthermore, in both the three and two-dimensional phase space trajectories a more distinct coactivation and deactivation of brain networks was observed in the local condition compared to the global condition. Similarly, when inspecting the phase space trajectories of two of the three brain networks against each other in all three possible combinations, younger adults exhibited a more sharply defined time specific coactivation and deactivation of singular brain networks compared to slower and less distinct transitions observed among older adults (Fig. S3).

**Fig. 3.**
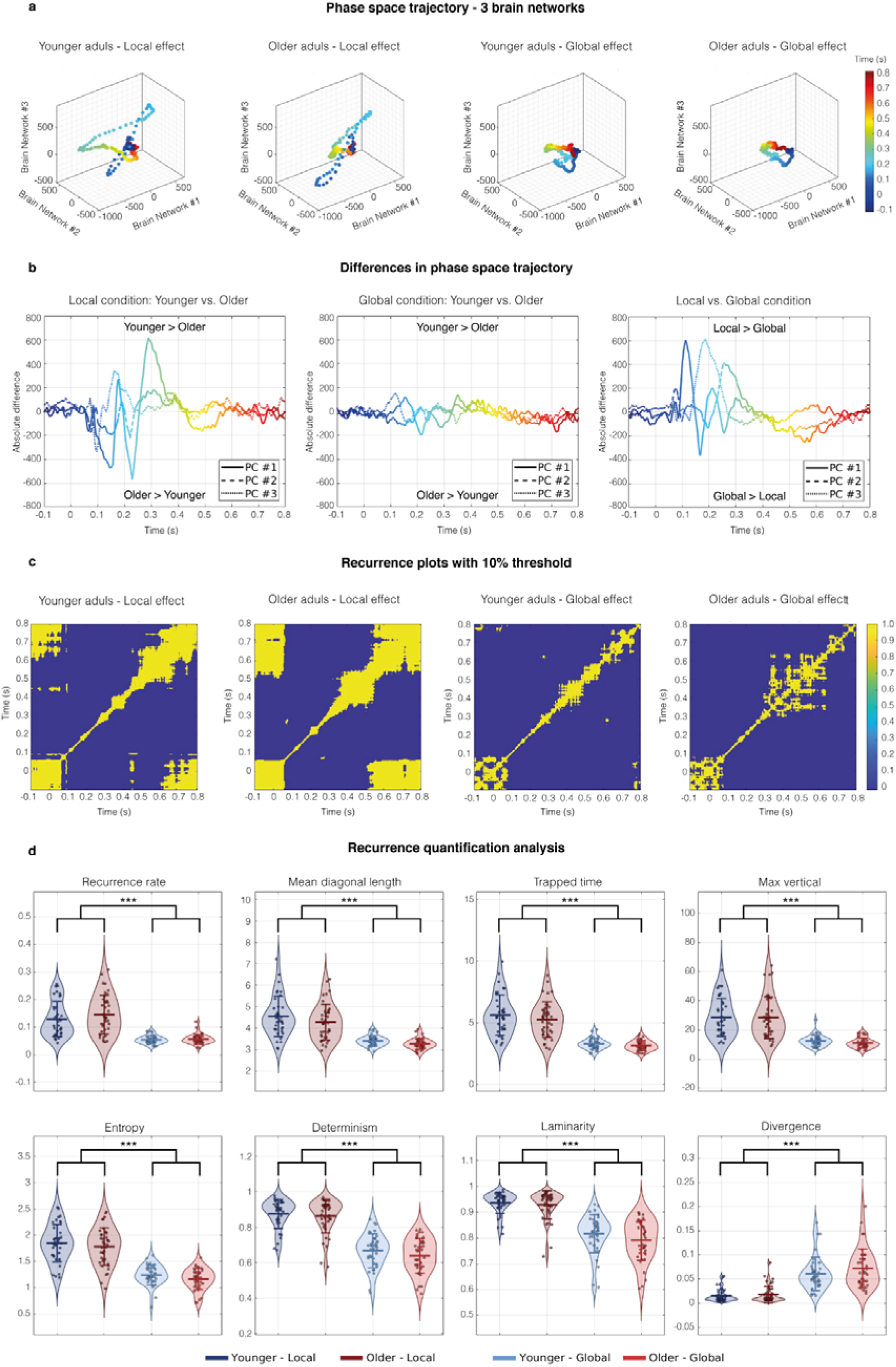
| Phase space embedding and recurrence quantification analysis of BROAD-NESS-derived brain network dynamics. **a**, Phase space trajectories averaged across participants (n = 77) for each experimental condition and age group, obtained by embedding the time series of the three primary BROAD-NESS networks into a three-dimensional state space. Each dot represents the state of the brain at a given time point, colour-coded by time (in seconds after the onset of the fifth tone). For specific values at each time point, see Table S5. **b**, Plots showing differences in three-dimensional phase space trajectories between age groups in each condition (left, centre) and differences between the conditions across age groups (right). **c**, Recurrence plots computed from the phase space trajectories shown in a) are averaged across participants, depicting pairwise Euclidean distances between time points above and below a 10 % threshold. Blue indicates larger distances (i.e., less similar brain states) that fall below the 10 % threshold, while yellow indicate greater similarity and temporal recurrence above the 10 % threshold. Fig. S4 depicts the unthresholded recurrence plots. **d**, Violin plots of eight essential recurrence quantification analysis metrics. The violin plots display the smoothed distribution of the data, along with the individual values, the group mean, and bars representing one standard deviation. The plots illustrate the statistically significant comparison between the local and global condition. These metrics quantify the predictability, complexity, and temporal persistence of network state trajectories. Significance levels: * p < .05, ** p < .01, *** p < .001; all p-values were significant after false discovery rate (FDR) correction.

When inspecting the thresholded recurrence plots derived from these trajectories (Fig. 3c), there was clear tendency for larger recurrency in the local condition across groups. Recurrency here refers to how often a previously occupied state is revisited reflecting how often the system’s dynamics return to similar configurations over time. For eight central recurrence metrics (such as recurrence rate, trapped time and entropy), effects of age group, condition and interaction was examined. All metrics showed significant condition effects (all FDR corrected p-values ≤ 1.6 × 10LL), with no significant main effects of group and no significant group × condition interactions (all FDR corrected p-values ≥ .16). Specifically, all metrics except divergence were increased in the local compared to the global condition, whereas divergence was increased in the global condition. These results indicate that, relative to global pattern processing, the local deviance condition is associated with more recurrent, predictable and temporally persistent network trajectories, while global processing is characterised by faster separation of trajectories, less stable, and more exploratory dynamics.

### Spatial gradient embedding of BROAD-NESS-derived network topographies

To further characterise the spatial organisation of the BROAD-NESS-derived networks, we applied a clustering approach to the voxel-wise activation maps of the first three principal components underlying the three main brain networks. Each voxel was embedded in a three-dimensional space according to the contribution of the voxel to each brain network (thresholded at mean ± 1 standard deviation), enabling a compact spatial representation of network co-involvement across the brain. Clustering analyses using k-means (performed on z-scored values and repeated 100 times per *k*, ranging from two to 40) revealed that a 16-cluster solution provided the most robust and reproducible partitioning of voxels in this space, as evaluated using silhouette coefficients. To ensure the reliability of this clustering solution, we repeated the entire clustering and silhouette evaluation procedure 5,000 times and consistently found that *k* = 16 yielded the highest silhouette scores across most repetitions. This optimal solution is visualised in Fig. 4, which shows the identified clusters projected back onto anatomical brain templates, while detailed information on the brain voxels forming these sixteen clusters is reported in Table S6, S7 and S8. For full transparency, Fig. S5 display clustering results and silhouette coefficients across the full range of *k* values, highlighting the robustness of the solution and the consistency of the results across adjacent clustering solutions. The sixteen-cluster structure provided a clear and interpretable mapping of how different voxel populations contributed to the three dominant networks. This analysis confirmed the presence of voxels strongly engaged in two of the three main brain networks, particularly within the auditory, cingulate and frontal cortices. Two examples are given: (i) Cluster #2 identified a group of voxels that contribute to the common auditory processes involved in Brain Network #1 and #2 within the left Heschl’s gyrus, Rolandic Operculum and Insula. (ii) Cluster #14 isolated a group of voxels that exclusively contributed to the frontal regions involved in Brain Network #2 and #3 within bilateral Rectus, and the right Olfactory Bulb and Subgenual Anterior Cingulate Cortex. We also observed voxels selectively contributing to only one network. For instance, bilateral Precuneus and the Posterior and Medial Cingulate cortices in Brain Network #1 (e.g. Cluster #4, #12 and #16), the left auditory cortex including Heschl’s gyrus in Brain Network #2 (Cluster #6 and #11) and the right auditory cortex including the Heschl’s gyrus in Brain Network #3 (Cluster #9 and #10). Finally, the analysis revealed a spatially coherent cluster of voxels with minimal or no contribution to either network, likely representing non-task-specific background activity (Cluster #1). Voxels within this cluster covered large parts of the frontal and occipital lobe and medial regions minimally implicated in the local-global paradigm.

**Fig. 4.**
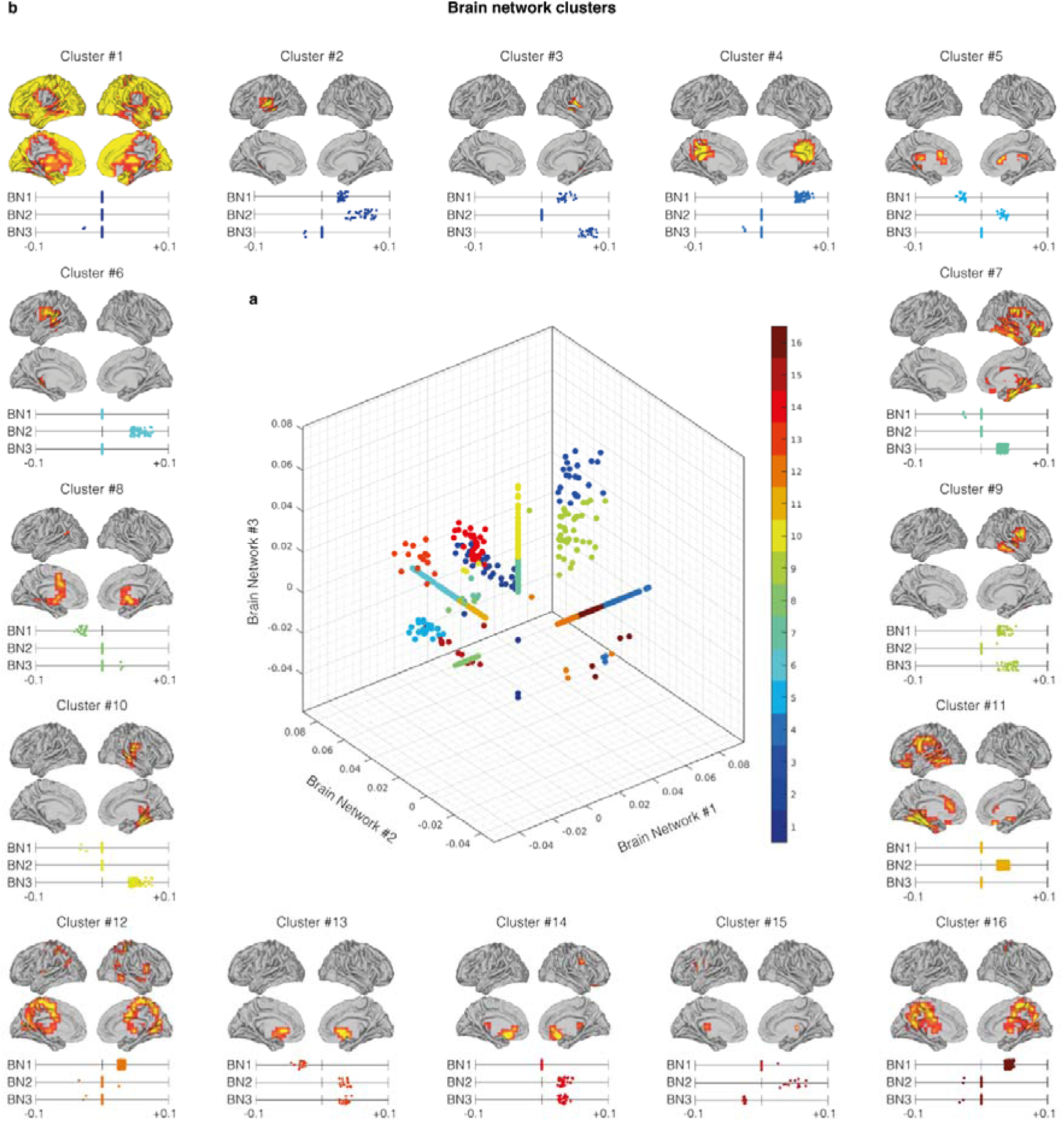
| Spatial gradient embedding and voxel clustering analysis. This figure illustrates the analysis of voxel-wise spatial activation patterns derived from the three main BROAD-NESS brain networks. **a**, Three-dimensional (3D) scatter plot showing the distribution of all brain voxels according to their spatial activation values in the brain networks obtained from the first three principal components revealing the underlying network involvement. The coloured voxels reveal the spatial gradients of the sixteen clusters identified by the optimal clustering solution, highlighting distinct patterns of co-involvement across the three networks. **b**, Brain template projections of the sixteen clusters from the optimal solution and their corresponding activation coefficients on each of the three main brain networks. These maps illustrate the spatial distribution of voxel groups identified in the central scatter plot revealing coherent and anatomically interpretable clusters, including shared, selective, and opposing contributors to the three BROAD-NESS networks. See Fig. S6 for a histogram of silhouette coefficients across 5,000 clustering iterations for each tested solution (k = 2 to 40). The peak at k = 16 confirms the optimal number of spatial clusters, based on maximum silhouette coefficient consistency.

### Brain network modularity analysis

To assess whether the large-scale brain networks engaged by the local–global paradigm were more constrained or more broadly distributed, we applied BROAD-NESS at the individual level and derived effective dimensionality from the eigenspectrum of the resulting covariance matrices. ED quantifies the effective number of orthogonal components contributing to the signal, with higher values indicating broader and less constrained network involvement.

Across all groups and conditions, the cumulative variance explained increased steeply for the first principal components before gradually approaching an asymptote (Fig. 5a), indicating that a limited subset of components accounted for the dominant whole-brain signal. However, the rise in explained variance was consistently delayed in the global compared to the local condition. This pattern suggests that processing global stimulus regularities involved a larger set of orthogonal network components.

**Fig. 5.**
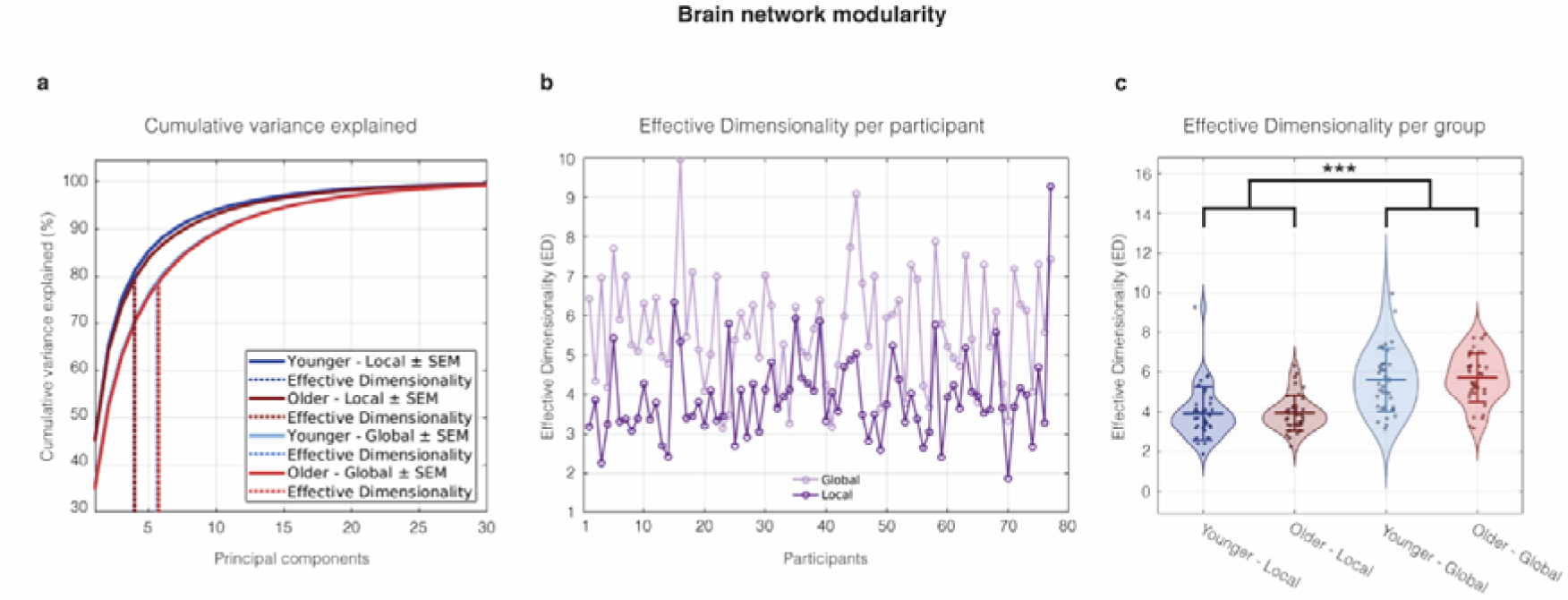
| Brain network modularity analysis. **a**, Cumulative variance explained by principal components when BROAD-NESS is applied at the individual level aggregated for groups and conditions in separation. Shaded traces represent the standard error of the mean. Vertical dashed lines indicate the number of components corresponding to the effective dimensionality of each group in each condition. **b**, Effective dimensionality values for each participant, plotted separately for the local and global condition. Each data point reflects data at participant level. **c**, Group-level distributions of effective dimensionality for younger and older participants, split by local and global networks. The violin plots display the smoothed distribution of the data, along with the individual values, the group mean, and bars representing one standard deviation. No significant group or interaction differences were observed. Significance levels: * p < .05, ** p < .01, *** p < .001; The p-value was significant after Bonferroni correction.

At the individual level, ED varied substantially across participants (Fig. 5b). Nevertheless, ED was higher in the global condition (M = 5.67, SD = 1.40) than in the local condition (M = 3.94, SD = 1.12), with 71 out of 77 participants showing this pattern. The within-subject difference (global − local) showed a mean of 1.74 (SD = 1.37).

A 2 × 2 mixed ANOVA with Condition (Local, Global) as a within-subject factor and Group (Younger, Older) as a between-subject factor revealed a robust main effect of Condition,

F(1,75) = 121.34, p < .001, partial η² = .62, indicating substantially higher ED in the global condition. The effect size was large, suggesting that approximately 62 % of the explainable within-subject variance was attributable to condition differences. There was no evidence for a main effect of Group, F(1,75) = 0.08, p = .779, partial η² = .001, nor for a Group × Condition interaction, F(1,75) = 0.08, p = .780, partial η² = .001. Thus, the increase in ED from local to global conditions was comparable between younger and older adults (Fig. 5c). Condition effects remained significant after Bonferroni correction (p < 0.001), while Group and Group x Condition effects remained insignificant (p = 1 in both cases). Although Anderson–Darling tests indicated deviations from normality in the local condition within each age group, the mixed ANOVA is generally robust to moderate non-normality in balanced designs. Moreover, the effect of Condition was highly consistent across participants, supporting the stability of the observed effect.

Together, these findings indicate that the local condition engaged more compact and constrained network configurations, whereas the global condition required a broader and more distributed set of network components to explain the variance of the data. This pattern is consistent with the idea that the processing of global deviations recruits more widely distributed large-scale connectivity patterns relative to the processing of local deviations, although these analyses alone cannot determine whether these differences arise from changes in network strength, complexity, noise structure, or a combination of factors.

### Investigating BROAD-NESS networks in group and condition separated data

To evaluate whether the time series and activation patterns identified by the BROAD-NESS methodology was robust to other grouping and averaging choices, we repeated the PCA decomposition on activation matrices generated separately for each age group in each condition. This procedure yielded one activation matrix per group-condition combination, allowing us to examine eigenspectra, time series, and spatial network structure under more constrained data partitions providing further insights of potential group and condition related differences. Across all datasets, the eigenspectra (Fig. 6A) showed that the first few principal components accounted for the majority of variance in the data, with sharp drops after the first three components. This pattern suggests that a relatively low-dimensional set of network processes captures most of the signal structure, even when PCA is performed on group– and condition-separated data.

**Fig. 6.**
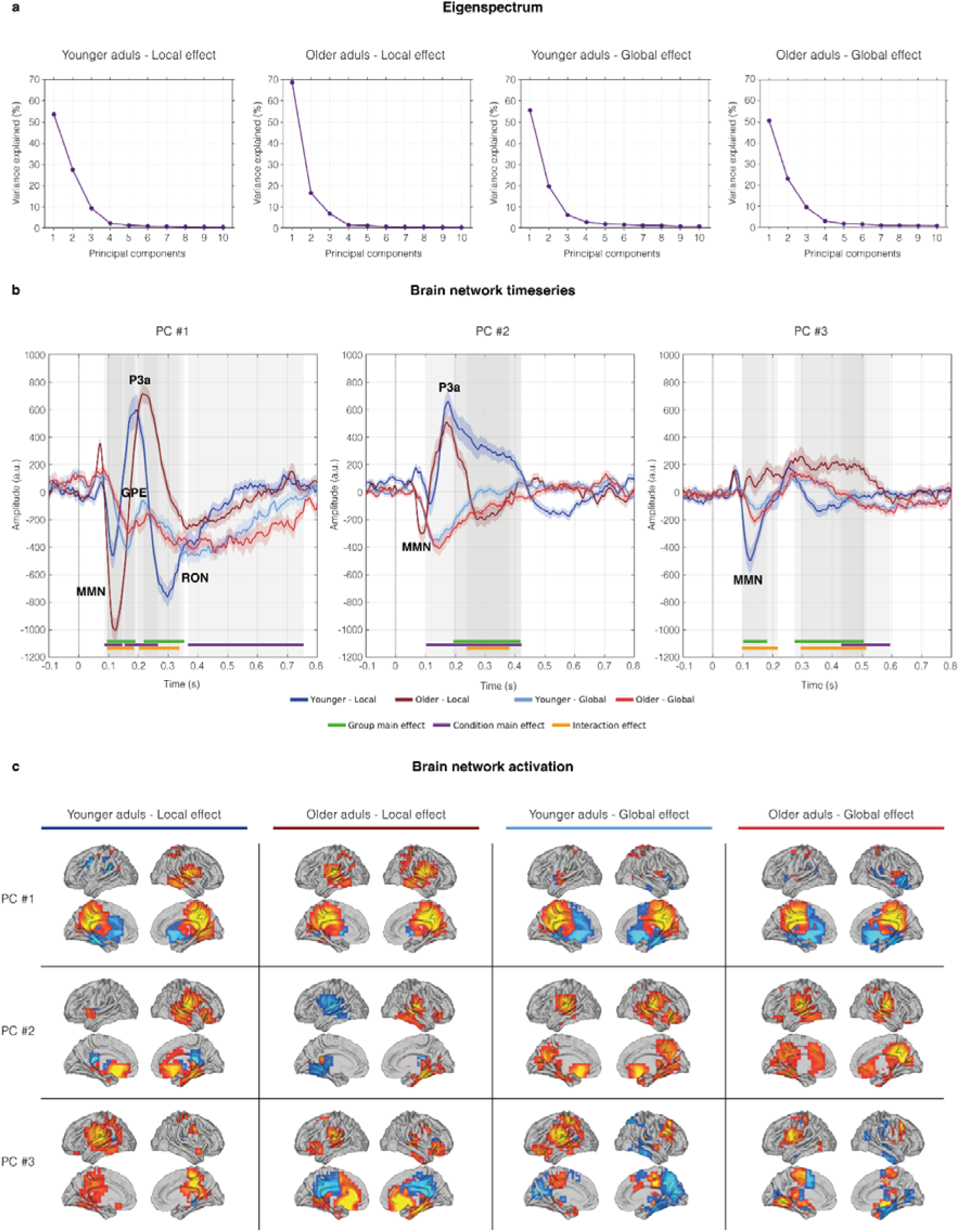
| Time series and brain network activation patterns when BROAD-NESS is applied to group and condition separated data. The figure illustrates the eigenspectra, time series and spatial activation patterns of brain networks when the BROAD-NESS framework is applied to group and condition separated data, which provides unique activation matrices for each group-condition separated dataset. **a**, Eigenspectra for group and condition separated data. Each plot displays the variance explained by the first 10 principal components. Across all conditions, the first three components account for most of the variance, indicating that only a small number of components capture most of the signal structure. **b**, Time series of the brain networks obtained from first three principal components (PC #1–PC #3) from group and condition separated data. Shaded areas indicate standard errors of the mean. Significant group, condition and interaction effects are indicated by green, purple and yellow lines. Time = 0 s. indicates the onset of the fifth tone in the local-global paradigm. **c**, Spatial activation maps of the brain networks obtained from the three first principal components for group and condition separated data. The brain templates illustrate the spatial extent of the networks, with yellow voxels contributing the most and light blue voxels contributing the least to the time series. Only voxels with values exceeding the mean by more than one standard deviation in absolute terms are depicted.

Inspection of the time series from the brain networks identified by the first three components revealed systematic age– and condition-related modulations (Fig. 6B). Brain Network #1 showed that tendencies in group, condition and interaction effects remained the same as in the main analysis revealing pronounced early local responses in older adults (MMN, P3a) and an enhanced later local RON response among younger adults. Brain Network #2 showed stronger local P3a responses in younger adults relative to older adults, demonstrating the same pattern observed in the main analysis. Brain Network #3 reflected increased MMN activity followed by later negative deflections, again with clearer expression in the local condition. Significant main effects of group and condition, as well as their interactions, occurred primarily within windows corresponding to ERF responses, and were more frequent in the local condition, suggesting a condition-specific selectivity in network engagement. See Table S9 for specific results.

Spatial maps of the first three components further highlighted meaningful differences across groups and conditions (Fig. 6C). Brain Network #1 involved bilateral coactivation of Medial and Posterior Cingulate cortices, Precuneus and Parietal areas, closely resembling the results of the main analysis. Interestingly, this coactivation differed among age-groups in the local condition, as older adults showed connections to the auditory cortices of both hemispheres, while younger adults predominantly utilised the right hemisphere. Brain Network #2 in the local condition captured activation of the right Caudate, Insula, Rolandic Operculum and Fusiform gyrus in both groups, while younger adults showed a stronger frontal engagement of the right Rectus, Anterior Cingulate Cortex and Olfactory Bulb. In the global condition, younger adults activated similar regions as in the local condition, while these regions were also connected to bilateral Precuneus activation. A similar pattern was observed for older adults, that showed a brain network connecting the auditory cortices to both frontal ACC and Rectus while also activating Precuneus and Medial/Posterior Cingulate Cortices. In Brain Network #3, younger adults displayed activation of left temporo-central areas including pre and post central areas, Cingulate cortices and the left temporal lobe. Older adults displayed activation of left Insula and Caudate areas across conditions and a very pronounced frontal activation in the local condition connecting bilateral Rectus, ACC and Olfactory Bulb areas to Caudate and Fusiform areas. See Table S10 and S11 for MNI coordinates and voxel counts within each brain area.

In sum, these analyses demonstrate that the time series and neural activation of the BROAD-NESS networks remain largely robust when PCA is applied to group– and condition-specific data. Though general tendencies in ERF responses and overall brain network structures remain consistent with the main analysis, this analysis also shows nuances in specific group and condition related differences of the time series and underlying brain network activation.

## Discussion

### Summary

In the present study, we applied BROAD-NESS, a PCA-based broadband network decomposition framework, to source-reconstructed MEG data acquired from the auditory local-global paradigm, where both sensory and pattern violations occur. Our two primary aims were to characterise hierarchical prediction error signalling within large-scale brain networks and its multivariate embedding in phase space and examine how these processes are altered in healthy aging. Three dominant orthogonal brain networks obtained from the first three principal components accounted for 91.5 % of variance in the data and were robustly recoverable across alternative PCA implementation strategies. Three key findings emerged in the study. First, automatically elicited prediction error signalling towards sensory deviancy was increased in aging and relied on a primary network connecting bilateral auditory regions to Medial and Posterior Cingulate Cortices strongly associated with the recruitment of attentional neural resources. Conversely, neural responses indexing the reorientation of attention following sensory violations and prediction error responses from global pattern violations were attenuated with age. Second, the processing of local violations involved a more recurrent and deterministic recruitment of the three main brain networks across time, while the processing of global violations was more divergent and exploratory. Third, the processing of global pattern violations was characterised by higher effective dimensionality of principal components. This suggested involvement of a wider range of orthogonal brain networks across the brain, compared to local violation processing which was more confined to fewer brain networks.

Together, these findings nuance existing predictive coding models of aging, where predictive processes indexed by ERF responses are expected to uniformly attenuate in strength independent of task demands. Instead, the present findings suggest that that these processes can enhance with age, that they are rooted in a large attention-related cingulo-parietal network and that dynamical distinctions between local and global processing are preserved with age. Below, we discuss each major set of results in turn, situating them within predictive coding and aging theoretical frameworks, while also considering alternative interpretations and methodological limitations.

### Large-scale network organisation of hierarchical prediction errors

We applied BROAD-NESS to source-reconstructed MEG data, and revealed three significant PCs that accounted for 51.78 %, 24.57 %, and 15.12 % of the variance in the data and displayed distinct network characteristics. Brain Network #1 linked bilateral auditory cortices including Heschl’s gyri, to Medial and Posterior Cingulate Cortices and Precuneus areas. These Parietal regions have repeatedly been implicated in attentional monitoring, salience regulation, and integrative cognitive control^51^. In contrast, Brain Network #2 and #3 were characterised by a pronounced involvement of the left and right auditory cortices and bilateral frontal regions including ACC and Rectus. This organisation suggests that the auditory cortex participates in multiple concurrent network configurations. In one configuration the auditory cortices are connected to cingula-parietal and are likely involved in both the detection of violations and in the updating of internal models of the environment – as both MMN and P3a were prominent within this network. In another configuration, the auditory regions of each hemisphere are connected to frontal areas including anterior cingulate circuits, which might suggest that they provide support for updating of internal predictive models. This is because as only P3a was prominently elicited within these networks, which aligns with previous studies, reporting that the ACC is one of the neural generators of the P3a response^34,41^.

Interestingly, these findings reveal both convergences and divergences with recent work on conscious predictive processes during working memory^52^ and long-term memory encoding^53^ and recognition of auditory sequences^9,54–57^. Similar large-scale networks have been reported, involving auditory cortices, medial and anterior cingulate regions, hippocampus, insula, and fronto-opercular areas. However, in contrast to those studies, the present results indicate a functional dissociation between left and right hemispheres, as reflected in the separation of Networks #2 and #3. Such hemispheric differentiation was not observed in studies of conscious recognition, which instead showed a higher relative reliance on medial cingulate gyrus^48^. This pattern suggests that while a common set of regions underpins predictive processing across conscious and pre-attentive contexts, their network configuration may differ depending on cognitive demands, particularly with respect to hemispheric specialisation and relative involvement of the medial cingulate.

Finally, to further characterise the spatial organisation of these networks, we embedded voxel-wise activation coefficients on each of the three brain networks within a three-dimensional gradient space and applied k-means clustering. A 16-cluster solution revealed spatially coherent and anatomically interpretable voxel groups. While several clusters contributed positively to one or two brain networks, no cluster showed contributions to all three networks. This illustrates how the three BROAD-NESS networks capture distinct but partially overlapping patterns of brain network co-involvement, embedded in continuous gradients rather than sharply segregated modules.

### Aging-related hierarchical alterations in prediction error signalling

Our primary aging-related finding concerned the pronounced alterations in ERF responses observed with age and the underlying brain networks supporting these processes. Older adults exhibited enhanced MMN– and P3a responses within an auditory-cingula-parietal network. This indicated an increased automatically elicited prediction error signalling towards sensory violations. More specifically, this indicated an increase in the processes involving the detection of deviancy and the updating internal models of sensory information. Conversely, more cognitively demanding ERF responses such as the late local RON response and the global prediction error response showed age-related attenuation within this network. Together, this pattern is consistent with results from a recent EEG study utilising the local-global paradigm, where older adults show no differences in MMN and P3a while the global prediction error response and RON responses were attenuated and predictive of future working memory decline^21,58^. The global prediction error response is widely considered to index conscious context updating and working-memory revision, with generators spanning parietal, cingulate, and frontoparietal networks^34,59^. Age-related reductions in the amplitude of the global prediction error response are therefore commonly attributed to impairments in higher-order contextual integration and working-memory updating^21,32^. Similarly, the RON response involves cognitively demanding processes to reorient attention towards the paradigm after distractions from sensory deviancy. This suggests an age-related difference in the recruitment of neural resources during complex tasks, where sensory-based are being prioritized ahead of more cognitively demanding processes. Furthermore, in a complex auditory memory paradigm involving the same participants as in the present study, an age-related enhancement of both an early negative response and later, slower positive components was observed^60^. This pattern was interpreted as reflecting a compensatory mechanism involving increased recruitment of attentional resources, which may contribute to the partial preservation of auditory and general memory abilities observed in this and similar cohorts^61,62^. Taken together, these results suggest that in complex tasks, age-related enhancements of sensory deviancy detection may be enhanced through attention-related Medial and Posterior Cingulate brain network involvement. At face value, this pattern challenges the canonical scalp-level literature predominantly reporting an age-related attenuation of prediction error signalling towards sensory violations^18,63,64^. Rather than a uniform reduction, our results suggest that aging involves a reallocation of attention-related neural resources, enabling enhancements of automatic, sensory-based prediction error responses – potentially at the cost of more cognitive processes involved in conscious working memory updating and reorientation of attention towards the paradigm. This pattern is consistent with the notion that older adults remain sensitive to sensory violations but show reduced capacity for sustained attentional reorientation and contextual revision, as these processes strongly dependent on working memory functions^41,58^.

In addition, these processes may be supported by different network configurations in aging. When specifically examining age-related differences when applying BROAD-NESS to group and condition separated data, the primary brain network (PC #1) showed connections between bilateral auditory cortices and cingula-parietal regions in older adults – whereas younger adults exclusively displayed connections between the right auditory cortex and cingula-parietal regions. This supports the Hemispheric Asymmetry Reduction in OLDer adults model (HAROLD), where aging is theorized to reduce lateralisation in favour of a bilateral recruitment of networks across hemispheres^65^. This suggests an age-related reduction in the hemispheric specialisation of predictive processing circuits, where bilateral auditory cortices rather than the right auditory cortex are connected with central Cingulate cortices to process sensory violations.

### Stimulus-specific adaptation as an alternative explanation

A critical interpretive challenge concerns the potential contribution of stimulus-specific adaptation (SSA) to age-related differences in early deviance responses. SSA is ubiquitous in the auditory cortex and contributes to MMN generation alongside predictive mechanisms^66,67^. Importantly, SSA is known to decline with age^68,69^, raising the possibility that group differences in MMN amplitude may partly reflect decreased neural adaptation rather than enhanced prediction error signalling. The local-global paradigm is inherently repetitive, which increases susceptibility to adaptation confounds. If older adults exhibit reduced adaptation to deviant tones, MMN-like difference waves may be amplified even in the absence of enhanced prediction error computation. This issue has been highlighted in critiques of the paradigm’s reliability in eliciting canonical MMN responses^70^, while in recent work a limited correspondence between MMN measures derived from oddball versus local-global designs has been shown^71^. Thus, the observed age-related enhancement of early components within Brain Network #1 could reflect adaptation-related variance becoming more globally distributed rather than compensatory recruitment of attention-related neural cortices. However, SSA effects are not deemed likely beyond MMN for two central reasons: (i) An age-related enhancement of P3a is observed, which is not known to be modulated by SSA. (ii) The dominant involvement of cingula-parietal regions in Brain Network #1, where SSA is not known to be present suggests involvement of attention-related neural resources. Despite these counterarguments, future work should incorporate paradigms designed to minimise SSA including e.g. roving standards or increased stimulus variability to disentangle adaptation from predictive coding contributions.

### Dynamical regimes depend on the hierarchical level of prediction error signalling

Beyond analyses of ERF responses, BROAD-NESS enabled a dynamical systems characterisation of brain network trajectories. Phase-space embeddings of network time series revealed that local deviants elicited comprised, recurrent trajectories, whereas global deviants induced more dispersed trajectories. Recurrence quantification analysis confirmed robust condition effects across all included metrics, with local processing showing higher recurrence, determinism, trapping time, diagonal length, max vertical, laminarity, and entropy, while global processing showed higher divergence. These findings are theoretically consistent with hierarchical predictive coding accounts. Local deviants violate short-lived sensory regularities and can be processed through relatively confined, recurrent circuits that give rise to easily identifiable and pronounced ERF responses. Global deviants require integration over longer time scales, revision of higher-order predictions, and engagement of conscious access mechanisms associated with the activation of global workspace systems across wide-ranging brain networks within the cortex^7,45,72–76^. Furthermore, ERF responses elicited towards global pattern violations are less pronounced and have a higher latency^7^. Importantly, these dynamical distinctions were largely age-invariant, suggesting that aging remodels the spatial embedding of deviance responses but does not abolish the qualitative dynamical separation of sensory and contextual prediction error signalling.

### A larger network organization observed in global deviance processing

Similarly, ED analysis provided converging evidence that the processing of global pattern violations engages a broader network recruitment across the brain. The processing of global deviants required a significantly increased ED across participants, indicating that more orthogonal components were required to account for signal variance. This aligns with the consistent findings that contextual violations recruit more distributed integrative systems consistent with global workspace theories and the conscious working memory dependent mechanisms involved in processing pattern violations^7,72^. However, ED is ultimately a covariance-based index rather than a direct measure of brain network complexity. Higher ED could reflect greater engagement of task-relevant networks but also increased neural noise or strategy heterogeneity. Thus, we interpret the ED increase as evidence for broader variance distribution during global processing without equating it unambiguously with more sophisticated computation.

### Limitations and future directions

Three limitations should be acknowledged. First, the study was cross-sectional, preventing direct inference about within-individual aging trajectories. Second, SSA remains a plausible confound for early component differences, motivating future meticulous manipulations of the stimuli incorporated within local-global paradigms. Third, frequency-specific analyses may provide enhanced mechanistic traction, given predictive coding theories linking distinct oscillatory bands to top-down and bottom-up signalling^2,77^. Future work should therefore aim to address these aspects to further advance knowledge on age-related changes in hierarchical prediction error signalling.

## Conclusion

In summary, aging does not simply attenuate hierarchical prediction error signalling. Rather, in complex tasks, the aging brain is capable of enhancing sensory-based processing of individual stimuli at the expense of more cognitively demanding processes involving conscious working memory and the reorientation of attention. By applying the PCA-based BROAD-NESS methodology to source-reconstructed MEG data from the auditory local–global paradigm, we identified three concurrent whole-brain networks supporting predictive processing. Aging was associated with an enhancement of local prediction error signalling within a primary brain network connecting cingula-parietal areas to the auditory system of the temporal lobes. These attention-related enhancements were exclusive to early, sensory-based processes among older adults, while cognitive processes involved in the reorientation of attention and working memory functions were attenuated. Crucially, the dynamical distinction between local and global processing remained preserved, with global violations engaging broader, higher-dimensional network dynamics. These findings underscore the importance of whole-brain network approaches in refining predictive coding models of aging and elucidating dynamical aspects embedded in the processing of sensory and pattern violations.

## Methods

### Participants

The sample comprised 77 participants (43 females), that were distributed in two age groups: (i) A younger adult group comprising 37 participants (18 females) aged 18 to 25 years, (mean age: 21.89 ± 2.05 years); (ii) An older adult group comprising 40 participants (24 females) aged 60 to 81 years (mean age: 67.50 ± 5.46 years). All participants held Danish nationality. Inclusion criteria encompassed: (i) Overall good health with no reported neurological or psychiatric issues, (ii) chronological age of either 18 – 25 years or 60 years and above, (iii) normal hearing adjusted to age, (iv) normal or corrected-to-normal vision (e.g. via contact lenses), and (v) a clear understanding of and agreement to the provided participant information. Exclusion criteria encompassed: (i) use of prescribed medication affecting the central nervous system, (ii) neurological or psychiatric conditions, (iii) unwillingness to cooperate or verbally agree to participation, (iv) MRI contraindications, (v) being 26 to 59 years old, and (vi) impaired hearing. The Institutional Review Board of Aarhus University, Denmark, approved the project (case number: DNC-IRB-2021-012). All experimental procedures were in compliance with the Declaration of Helsinki – Ethical Principles for Medical Research (World Medical Association Declaration of Helsinki, 2013). All participants were informed about the purpose of the study before giving consent to participate. A brief demographic questionnaire was administered prior to MEG data collection, obtaining information regarding age, highest level of achieved education and biological sex. See table 1 for an overview.

**Table 1.** Demographic information. Data is reported in whole numbers or mean ± SD.

|  | Younger adults (n = 37) | Older adults (n = 40) | p values |
| --- | --- | --- | --- |
| Sex (females/males) | 18 / 19 | 24 / 16 | - |
| Age in years | $21.89 \pm 2.05$ | $67.50 \pm 5.46$ | $< 0.001^{***}$ |
| Years of education | $13.84 \pm 1.85$ | $14.88 \pm 3.07$ | 0.074 |

### Experimental paradigm

An adapted version of the auditory local-global paradigm was used^7^. The task consisted of 160 trials per block for a total of 4 blocks. Each block consisted of a combination of patterns consisting of either five repetitive tones, or patterns where the fifth tone deviated in tone frequency (Fig. S1). Two types of deviating fifth tones were used – an experimental deviating tone to induce a local sensory violation, and a deviating target tone to ensure continued attention towards the paradigm, as participants were instructed to press a button upon hearing this tone.

The local-global paradigm was designed to elicit neural responses at two hierarchical levels: Canonical local neural responses were expected to be elicited from sensory violations at the individual tone level, while canonical global neural responses were expected to be elicited from violations at pattern level. Both congruent and incongruent blocks were used (Fig. S1). In congruent blocks, local standard trials with five repetitive tones occurred 70 % of the time, while local deviant trials with a fifth deviating experimental tone occurred 23.75 % of the time. In both congruent and incongruent blocks, target tones were elicited 6.25 % of the time to ensure continued attention towards the paradigm. In incongruent blocks, local standard trials switched probability of occurrence with local deviant trials to create a global violation regarding the frequency with which stimuli was presented. The ordering of congruent and incongruent blocks was randomized among participants. The duration of each of the five tones in each trial was 80 ms followed by 20 ms of silence after each tone yielding a stimulus onset asynchrony of 100 ms within patterns. Between the five tone patterns the intertrial interval was 500 ms. Three pitch levels were used, centred around 440 Hz and spaced in semitone steps (−4, 0, +10 semitones). The local-global paradigm concluded after all 4 blocks lasting 10 minutes and 40 seconds excluding pauses, while the entire MEG session concluded after approximately 1 hour, as other tasks were also administered. The order of tasks in a session was randomized across participants. The global-local-paradigm was presented using Psychopy version 2023.2.3^78^.

Instructions were displayed visually on a screen at the beginning of each block, prompting participants to relax before starting and to look at a fixation cross presented at the centre of the screen. Instructions prompted participants to identify the target tone in each block by pressing a button on a joystick. A practice phase familiarized participants with detecting the target tone. Practice continued until participants produced three correct responses within 2 s of target onset within one practice round, ensuring task comprehension before continuing. Upon completing the practice phase, two congruent and two incongruent blocks were presented in random order. Auditory stimuli were delivered through a speaker placed 2 metres in front of the subject at a sound pressure level of ∼60 dB for 68 participants. For 9 participants sounds were presented at ∼70 dB, as they were 70 years or older and showed mild hearing impairment, typically associated with aging. Only auditory stimuli in the 125 – 650 Hz range were used, as these frequencies are minimally impacted by typical age-related hearing loss^79^. Event timing, response times, and task events were logged for offline analyses.

### Neural data acquisition

The MEG recordings were conducted in a magnetically shielded room at Aarhus University Hospital (AUH), Denmark, using an Elekta Neuromag TRIUX MEG scanner with 306 channels (Elekta Neuromag, Helsinki, Finland). Data was collected at a sampling rate of 1,000 Hz with an analogue filtering of 0.1 – 330 Hz. Prior to the recordings, participants’ head shapes and the positions of four Head Position Indicator (HPI) coils were registered relative to three anatomical landmarks using a 3D digitizer (Polhemus Fastrak, Colchester, VT, USA). This information was later used to co-register the MEG data with anatomical MRI scans. Throughout the MEG recording, the HPI coils continuously tracked the position of the head, allowing for movement correction during analysis. Additionally, two sets of bipolar electrodes were used to monitor cardiac rhythm and eye movements, facilitating the removal of electrocardiography (ECG) and electrooculography (EOG) artefacts in preprocessing.

MRI scans were obtained on a CE-approved 3T Siemens MRI scanner at AUH. The data included structural T1 (MPRAGE with fat saturation) scans, with a spatial resolution of 1.0 x 1.0 x 1.0 mm and the following sequence parameters: echo time (TE) of 2.61 ms, repetition time (TR) of 2,300 ms, a reconstructed matrix size of 256 x 256, an echo spacing of 7.6 ms, and a bandwidth of 290 Hz/Px. The MEG and MRI recordings were conducted on separate days.

### MEG data pre-processing

The raw MEG sensor data (comprising 204 planar gradiometers and 102 magnetometers) was initially pre-processed using MaxFilter^80^ (version 2.2.15) to reduce external interferences. We applied signal space separation (SSS) with the following MaxFilter parameters: downsampling from 1,000 Hz to 250 Hz, movement compensation via continuous HPI coils (default step size: 10 ms), and a correlation limit of 0.98 to reject overlapping signals between inner and outer subspaces during spatiotemporal SSS.

The MEG data was converted into Statistical Parametric Mapping (SPM) format and further processed and analysed using MATLAB (MathWorks, Natick, MA, USA), incorporating custom-built scripts (LBPD, https://github.com/leonardob92/LBPD-1.0.git) and the Oxford Centre for Human Brain Activity (OHBA) Software Library (OSL)^81^ (https://ohba-analysis.github.io/osl-docs/), which integrates Fieldtrip^82^, FSL^83^, and SPM^84^ toolboxes.

Next, the continuous MEG data underwent visual inspection via the OSLview tool to remove any large artifacts, where less than 0.1 % of the collected data was discarded. Independent Component Analysis (ICA) was then used (with OSL implementation) to eliminate eyeblink and heartbeat artifacts^85^. The ICA process involved decomposes the original signal into independent components and correlates these components with the activity recorded by the electrooculography (EOG) and electrocardiography (ECG) channels^86^. Components that showed a correlation at least three times higher than others were flagged as reflecting EOG or ECG activity. These flagged components were further validated through visual inspection, ensuring their topographic distribution matched typical eyeblink or heartbeat activity, and were subsequently discarded. The signal was then reconstructed using the remaining components. The signal was epoched using a 100 ms baseline before the onset of the fifth tone and 800 ms following this onset. To achieve an equivalent signal-to-noise-ratio between standards and deviants, an equivalent number of standard epochs were randomly selected. For each group and for each condition, signals from standard and deviant epochs were averaged in separate before being subtracted from each other.

### Source reconstruction

MEG is a powerful tool in detecting whole-brain activity with excellent temporal resolution. However, to fully understand the brain activity involved in complex cognitive tasks, it is essential to identify the spatial sources of this activity. This requires solving the inverse problem, as the MEG recordings reflect neural signals from outside the head but do not directly indicate which brain regions generated them. To address this, we employed beamforming algorithms^87–89^, using a combination of in-house-developed codes alongside the OSL, SPM, and FieldTrip toolboxes. The procedure consisted of two main steps: (i) Designing a forward model and (ii) computing the inverse solution.

First, a single-shell forward model was constructed using an 8-mm grid. This head model treats each brain source as an active dipole and describes how the activity of such a dipole would be detected across MEG sensors^90,91^. The MEG data was co-registered with individual T1-weighted MRI scans, using the 3D digitizer information to align the data. When individual anatomical scans were unavailable, an MNI152-T1 template with 8-mm spatial resolution was used to compute the forward model.

Second, a beamforming algorithm was applied as the inverse model. Beamforming uses a set of weights applied to different source locations (dipoles) to isolate the contribution of each brain source to the activity recorded by the MEG sensors^87,89,92^. This was done for every time point of the recorded brain data, allowing us to reconstruct the spatial sources of the MEG signal. The detailed steps for implementing the beamforming algorithm are provided below.

The data recorded by the MEG sensors (*B*) at time *t*, can be described by the following Eq. (1):

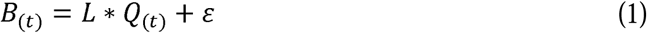

where *L* is the leadfield model, *Q* is the dipole matrix carrying the activity of each active dipole (*q*) at time *t*, and L is noise (for details, see Huang et al.^91^). To solve the inverse problem, *Q* must be estimated for each *q*. In the beamforming algorithm, a series of weights are computed to describe the transition from MEG sensors to the active dipole *q*, independently for each time point. This is reported in Eq. (2):

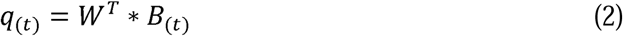

Here, the superscript *T* refers to transpose matrix. To compute the weights (*W*), matrix multiplication between *L* and the covariance matrix of MEG sensors (*C*) is performed. Importantly, the covariance matrix *C* was computed on the signal after concatenating the single trials of all experimental conditions. For each dipole *q*, the weights (*W_q_*) were computed as shown in Eq. (3):

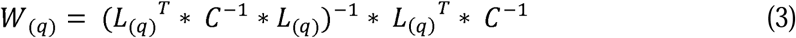

The computation of the leadfield model *L* was performed for the three main orientations of each dipole^90^. Before computing the weights, to simplify the beamforming output^93,94^, the orientations were reduced to one using the singular value decomposition algorithm on the matrix multiplication reported in Eq. (4):

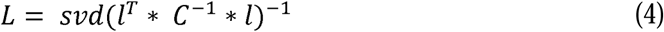

Here, *l* represents the leadfield model with the three orientations, while *L* is the resolved one-orientation model that was utilized in Eq. (3).

Once computed, the weights were normalised for counterbalancing the reconstruction bias towards the center of the head^87,95–97^ and applied to the neural activity averaged over trials, independently for each time point (Eq. 2) and experimental condition. This procedure returned a time series for each of the 3,559 brain sources^87,96^.

### Principal Component Analysis (PCA)

PCA was applied to the time series of the 3,559 reconstructed brain sources to disentangle broadband brain networks operating simultaneously. PCA is a dimensionality reduction method that transforms data into new variables, or principal components, which account for the most variance within the dataset. This is achieved by computing the eigenvectors and eigenvalues from the covariance matrix of the data and then projecting the data onto the directions of maximum variance. Traditionally, PCA simplifies high-dimensional data while retaining the most important information.

In this study, however, PCA was applied to a dense set of MEG-reconstructed brain data at the voxel level (3,559 brain voxel time series). The goal was not just to reduce the dimensionality but to identify brain networks obtained from PCs involved through the application of PCA on the 3,559 brain voxel time series.

In short, PCA operates as follows. First, the data is centred by subtracting the mean of each brain voxel timeseries from the dataset *X*, as represented by Eq. (5):

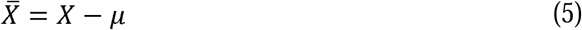

where µ is the mean vector of *X*.

Then, the covariance matrix *C* is computed on the centred data, as shown in Eq. (6):

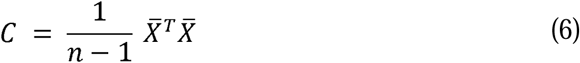

where *n* is the number of data points.

Then the eigenvalue equation for the covariance matrix is solved to find eigenvalues and eigenvectors, Eq. (7):

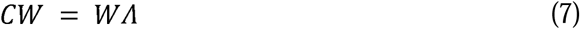

Where W is the eigenvector matrix (set of weights for each principal component) and A is the diagonal matrix of the corresponding eigenvalues (which indicate the amount of variance explained by each component).

Then, the eigenvectors w were used to compute the activation time series of each brain network *y* by multiplying them by the original data *X*, as shown in Eq. (8):

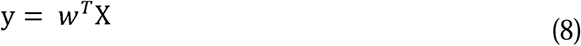

Furthermore, we computed the spatial projection of each component (spatial activation patterns *a*) in brain voxel space. This was done by multiplying the weights of the analysis (the eigenvectors w in this case) by the covariance matrix *C*, as shown in Eq. (9):

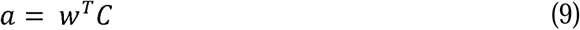

Interestingly, while previous studies on interpreting weights from multivariate analyses in MEG data have recommended calculating spatial activation patterns^98^, in this case, the relative contribution of each brain voxel to the network remains the same, whether using the direct eigenvector w or computing the spatial activation patterns.

It is important to note that PCA can be computed using different mathematical approaches that yield equivalent results. A commonly used alternative is to apply Singular Value Decomposition (SVD)^99^ directly to the mean-centred data matrix. This approach is often preferred in practice because it is more numerically stable and computationally efficient, especially for high-dimensional data (for instance, this is the solution currently implemented in the ‘pca’ function in MATLAB). However, since the SVD-based solution ultimately produces the same principal components and scores as the covariance-based eigendecomposition we describe above, we chose to retain the latter formulation. This choice is motivated by clarity and pedagogical value, as it offers a more intuitive explanation of the underlying principles.

Importantly, as detailed in the following section, the PCA procedure was applied across various scenarios to ensure a comprehensive evaluation of the algorithm’s performance. This included performing PCA in three different scenarios: (i) On data averaged across participants and conditions in the main analysis. (ii) On data from each participant in each condition in the brain network modularity analysis. (iii) On group and condition averaged data as a robustness-check and to elucidate more nuanced group– and condition related differences complementing the main analysis.

## Statistical analysis

After computing PCA on the group averaged data, we extracted the time series of the resulting brain networks for each participant in each condition (Local, Global)^48^. Followingly, we assessed group– and condition-related effects within a 2 × 2 mixed factorial framework, using separate permutation-based tests to evaluate the Group main effect, Condition main effect, and Group × Condition interaction. For each of the first three brain networks, we restricted the analysis to the 0 – 800 ms post-stimulus interval and performed cluster-based permutation tests over time following the framework of Maris and Oostenveld^100^. At each time point, we tested (i) a between-subject Group effect by comparing younger and older adults after averaging across conditions, (ii) a within-subject Condition effect by contrasting the global versus local condition across all participants, and (iii) a Group × Condition interaction defined as the difference in Global–Local contrasts between groups. Statistical inference was performed using cluster-based permutation tests over time with 5,000 permutations, where cluster-mass statistics controlled the family-wise error rate. Clusters of neighbouring supra-threshold time points were first defined using a pointwise cluster-forming threshold of p < 0.05. The resulting cluster masses (sum of absolute effect values) were then compared against a reference distribution of maximum cluster masses obtained under the null hypothesis by shuffling group labels for between-subject effects and applying random sign flips for the within-subject contrast. The family-wise error rate was controlled at α = 0.05 based on this max-cluster-mass distribution. For each PC and each effect, we obtained binary significance masks and corresponding time windows of significant clusters, which were then used to visualize coloured intervals indicating periods of significant group, condition, or interaction effects. All statistical analyses were implemented in MATLAB using the standardized routines described above and including the full sample of 77 participants.

### Phase space and recurrence quantification analysis of PCA-derived networks

To investigate the temporal dynamics and recurrence properties of the brain networks identified via the PCA-based BROAD-NESS methodology, we conducted a phase space and recurrence quantification analysis^101,102^ on the time series of the three main PCA components (brain networks). This approach allows for the examination of the brain’s timewise utilisation of the three main brain networks depicted in a phase space trajectory within a low-dimensional temporal space, revealing underlying temporal structures that may not be evident in the original signal.

We first constructed two– and three-dimensional phase space trajectories by treating the time series of the first three BROAD-NESS networks as independent axes in phase space. For each participant and experimental condition, the time-resolved trajectory in this space was computed. Each point in the resulting scatterplot represents the state of the brain at a given time point, embedded in the space defined by the three dominant networks. Next, recurrence plots (RPs) were calculated for each participant and condition. RPs reflect the degree to which the system re-enters similar network states over time^101,102^. Specifically, for each pair of time points, we computed the Euclidean distance between their respective phase space coordinates. Recurrence matrices were thresholded using a fixed ε-radius criterion (10% of the maximum distance) to obtain binary plots, where a value of 1 indicates a recurrence^101,102^.

Next, we quantified how these trajectories differ across age groups in each condition and across conditions by extracting the participant-averaged phase-space coordinates (PC #1 – PC #3). To examine differences in both negative and positive movements of phase space trajectories, we examined absolute differences, by comparing absolute differences in phase space coordinates at each time point for all three PCs, as shown in Eq. (10):

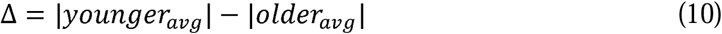

This was done when comparing age groups in the local and global condition to obtain averaged absolute time wise differences. This yielded positive values when younger adults exceeded older adults, and negative values when older adults exceeded younger adults.

Conversely, when comparing absolute differences between the local and global condition across age groups, we first computed group-averaged values of the time series of each of the three main brain networks within each condition, as shown in Eq. (11):

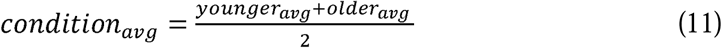

and then computed:

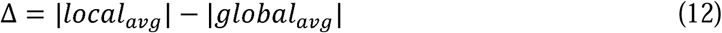

Following Eq. (12), we calculated recurrence plots (RPs) for each participant and condition. RPs reflect the degree to which the system re-enters similar network states over time^101,102^. Specifically, for each pair of time points, we computed the Euclidean distance between their respective phase space coordinates. Recurrence matrices were thresholded using a fixed ε-radius criterion (10 % of the maximum distance) to obtain binary plots, where a value of 1 indicates a recurrence^101,102^. From the thresholded RPs, we extracted eight standard metrics commonly used in nonlinear time-series and brain dynamics research. For full mathematical descriptions, see^102^.

1. Recurrence Rate (RR): proportion of recurrent points, indexing overall recurrency of the system.
2. Mean Diagonal Line Length (L): average length of diagonals, indicating temporal predictability.
3. Trapping Time (TT): average vertical line length, reflecting temporal persistence in a given state.
4. Maximal Laminar Duration (Vmax): length of the longest vertical line (i.e. period of state stability).
5. Entropy (ENTR): Shannon entropy of diagonal lengths, indexing complexity of the system’s evolution.
6. Determinism (DET): proportion of recurrence points forming diagonal structures, reflecting the stability of the system trajectories.
7. Laminarity (LAM): fraction of recurrence points forming vertical lines, reflecting intermittency.
8. Divergence (DIV): inverse of the longest diagonal line, linked to system sensitivity and divergence in state space.

These metrics were computed independently for each participant in each condition. Statistical analyses were performed using linear mixed-effects models implemented in MATLAB. For each metric, we fitted a model with Group (younger vs. older adults) as a between-subjects factor and Condition (global vs. local) as a within-subjects factor, as shown in Eq. (13):

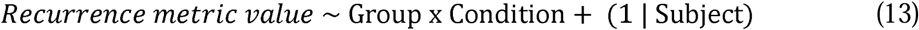

A random intercept for each subject accounted for repeated measures. Fixed effects were evaluated using F-tests with Satterthwaite-approximated degrees of freedom. For each metric, we extracted p-values for (i) the main effect of Group, (ii) the main effect of Condition, and (iii) the Group × Condition interaction. To control for multiple comparisons, all p-values across metrics and effects (Group, Condition, Interaction) were pooled and corrected using the Benjamini–Hochberg False Discovery Rate (FDR) procedure^103^. Reported p-values therefore reflect correction across the full family of tests. For visualization, violin plots were generated for each metric, showing the distribution of values for the four combinations of Group and Condition. Means and variability were displayed within each violin plot to aid interpretation.

### Spatial gradient embedding and clustering analysis of network topographies

To examine the spatial organisation of the dominant brain networks identified via the BROAD-NESS framework, we performed a spatial gradient embedding and clustering analysis on a voxel-wise activation map. This analysis was conducted on the data averaged across participants to obtain a robust, group-level representation of network topographies. We focused on the first three principal components (PC #1 – PC #3), as they represented the most important brain networks given calculations of effective dimensionality. For each component, we thresholded the voxel-wise spatial weights by setting values between mean ± 1 standard deviation to zero. This allowed us to retain only the most strongly contributing voxels. These thresholded values were then used to define a three-dimensional (3D) embedding space, where the position of each voxel was determined by its weight on PC #1 (x-axis), PC #2 (y-axis) and PC #3 (z-axis). This provided a gradient-like spatial representation, capturing how individual voxels contributed to the three brain networks. To identify spatially coherent voxel clusters within this 3D space, we applied k-means clustering^104^ to the embedded voxel coordinates. Prior to clustering, the data were z-scored to normalise both axes and ensure equal weighting of the three components. For each value of *k* (ranging from 2 to 40)^105^, we computed the k-means clustering in 100 repetitions with different random initialisations to minimise the effect of convergence to local minima^104^. To determine the optimal number of clusters, we computed the silhouette coefficient^106^ for each clustering solution. This metric assesses how well-separated the resulting clusters are by comparing the average distance between points within the same cluster to the average distance between points in neighbouring clusters. Higher silhouette values indicate better-defined, more distinct clusters.

However, due to the stochastic nature of k-means initialisation, the silhouette scores can vary slightly across runs. To account for this variability and ensure robust estimation of the optimal number of clusters, we computed silhouette scores 5,000 times. The final clustering solution was selected based on the number of clusters that was most frequently associated with the highest silhouette score across these repetitions. This method provided a robust and data-driven way to quantify the spatial gradients underlying the network topographies derived by the BROAD-NESSS method. Crucially, it enabled the identification of consistent and interpretable voxel clusters, including those that predominantly contributed to one network, those shared between two or three networks, and those with negative or minimal contributions. This approach offers a principled means to uncover the fine-grained spatial architecture of large-scale functional brain networks.

### Investigating BROAD-NESS networks in group and condition separated data

To enhance the robustness of our work and provide greater insight into the subtle differences associated with the computation of PCA on neural data, we examined a crucial aspect relevant to experimental settings. Specifically, when multiple participant groups and experimental conditions are involved, it is essential to determine whether it is more beneficial to compute PCA on group and condition separated data or on data aggregated across groups and conditions.

To address this, we computed both approaches and compared the results. Upon applying to PCA data averaged across conditions and groups in the main analysis, we computed PCA independently for each group in each condition. This allowed us to compare the brain networks and resulting time series computed using these two approaches and assess potential differences. In both approaches, the time series of brain networks were reconstructed independently for each participant using the weights of the activation matrix obtained through each method.

Specifically in the case of group and condition separated data, the original MEG data were organized as a 4D array with dimensions *sources × time × condition × participants* and were split into four subsets corresponding to younger-local, younger-global, older-local, and older-global data. This approach allowed us to evaluate whether the dimensional structure, temporal expression, and spatial organization of BROAD-NESS networks were preserved when network estimation was performed on more constrained group– and condition-specific datasets.

Participants were assigned to younger and older groups based on predefined aging criteria, and data were indexed to create four 4D matrices of size *sources × time × 1 × participants* providing one matrix per group–condition combination. These matrices were then used as input to the BROADNESS pipeline, which computes group-level network estimates while retaining participant-specific time series for subsequent statistical analyses.

Within each group–condition dataset, PCA was applied to the group-averaged MEG data to derive principal components capturing dominant patterns of covariation across brain sources over time. BROAD-NESS returns (i) an eigenspectrum depicting the variance explained by each component and (ii) the corresponding brain network time series for each participant. For each decomposition, we inspected the full eigenspectrum and quantified effective dimensionality from the eigenvalue distribution, but focused interpretation on the first three components, which consistently accounted for the largest portion of explained variance across datasets.

For each retained component, we examined (i) the associated temporal profile of the network time series and (ii) the spatial activation pattern across brain sources. Network time series were extracted and used to characterize event-related modulation. Spatial activation patterns were visualized by projecting component weights onto an 8 mm MNI152 template using the provided MNI voxel coordinates, yielding network topographies for each group-condition specific decomposition. This further allowed for a qualitative comparison of spatial network structures across groups and conditions.

To evaluate differences between groups and conditions in the temporal expression of the BROAD-NESS networks, we subjected the time courses of the first three components to timepoint-wise 2 × 2 analyses (reflecting groups and conditions) using cluster-based permutation testing. For each principal component, we contrasted group means (averaged over conditions), condition differences (global–local) and their interaction across the 0–800 ms time window, forming clusters of contiguous time points exceeding a cluster-forming threshold. Family-wise error was controlled via permutation-derived maximum cluster mass distributions.

The rationale of this analysis was: (i) To investigate whether effects observed in the main BROAD-NESS analysis were robust when network estimation was recomputed on group-and condition-specific data. (ii) To investigate potential nuanced differences in time series expressions and activation patterns, when applying BROAD-NESS at more specific partitions of the data.

### Brain network modularity analysis

To characterize the modular organization of large-scale brain network activity, we applied estimated temporally coherent large-scale network components using principal component analysis (PCA) on the covariance structure of data from each individual. For each participant and each condition, we obtained a sorted eigenspectrum depicting the proportion of variance explained by each principal component. This procedure was applied independently for all participants, producing one eigenspectrum per subject and condition.

To quantify the degree to which activity patterns were concentrated versus broadly distributed across components, we derived an estimate of effective dimensionality^50^ from the eigenspectrum.

Effective dimensionality was quantified using the participation ratio of the principal component eigenvalue spectrum. Following principal component analysis (PCA) of the BROADNESS brain network variance matrix, the vector of component variances A_i_ was extracted. Effective dimensionality (ED) was computed as:

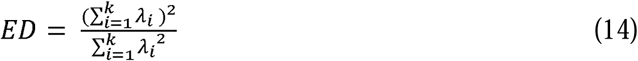

This measure reflects the number of components that effectively contribute to the variance structure, independent of the nominal dimensionality. Eigenvalues were constrained to be non-negative prior to computation. The measure is bounded between 1 and the total number of components.

Conceptually, higher ED values indicate that the total variance is distributed across a larger set of components, whereas lower values indicate that the variance is captured by a smaller, more compact network structure. ED was calculated for each participant in each condition. We visualized the distribution of ED values using scatter plots at the single-subject level and violin plots at the group level. Group differences were tested using a 2 × 2 mixed ANOVA with Group (younger, older) as a between-subject factor and Condition (local, global) as a within-subject factor. P-values were Bonferroni-corrected across the three main ANOVA effects.

To relate ED to the underlying eigenspectra, we also computed cumulative variance explained curves for each condition and group. For each participant, cumulative variance was calculated for the first 30 principal components. Group means and standard errors were derived separately for younger and older adults. Vertical reference lines were plotted at the group-level to illustrate how mean ED values corresponded to plotted curves indicating cumulative variance explained across principal components.

## Data availability

The pre-processed neuroimaging data generated in this study have been deposited in the Zenodo database and are publicly available: https://doi.org/10.5281/zenodo.18231641107

The code used to present the stimuli of the experimental paradigm used in this study has been deposited on GitHub and is publicly available: https://github.com/MathiasHoueAndersen/Hierarchical-Predictive-Processing/blob/main/Experimental_Paradigm.py

## Code availability

In-house-built code and functions used for the pre-processing of MEG data in this study are part of the LBPD repository which is available at the following link: https://github.com/leonardob92/LBPD-1.0.git

The BROAD-NESS toolbox is available at the following link and is required to run the data analysis pipeline: https://github.com/leonardob92/BROADNESS_MEG_AuditoryRecognition/tree/main/BROADNESS_Toolbox

The full data analysis pipeline used in this study is available at the following link: https://github.com/MathiasHoueAndersen/Predictive-Processing-In-Aging

## Acknowledgements

The Center for Music in the Brain (MIB) is funded by the Danish National Research Foundation (project number DNRF117).

L.B. is supported by Sapere Aude: Independent Research Fund Denmark (DFF) Research Leader (grant ID: 10.46540/5253-00003B), Lundbeck Foundation (Talent Prize 2022), Carlsberg Foundation (CF20-0239), Center for Music in the Brain, Linacre College of the University of Oxford and Nordic Mensa Fund.

M.L.K. is supported by Center for Music in the Brain and Centre for Eudaimonia and Human Flourishing, which is funded by the Pettit and Carlsberg Foundations.

## Author contributions statement

L.B., M.R. and M.H.A. conceived the hypotheses. L.B., G.F.R. and D.R.Q.M. designed the study. L.B., M.L.K. and P.V. recruited the resources for performing data collection and analysis. D.R.Q.M. designed the experimental paradigm used in the study. G.F.R., M.K. and L.B. collected the data. M.H.A. and L.B. performed pre-processing, statistical analysis and decomposition of the neural signal. L.B. and M.H.A. updated the code for the BROAD-NESS toolbox. M.L.K., P.V., H.R.S., K.M.L. and M.R. provided essential help to interpret and frame the results within the neuroscientific and analytical literature. M.H.A. wrote the first draft of the manuscript, which was primarily integrated by L.B. and M.R. The figures were prepared by M.H.A., L.B. and M.R. All the authors contributed to and approved the final version of the manuscript.

## References

1. Rao, R. P. N. & Ballard, D. H. Predictive coding in the visual cortex: A functional interpretation of some extra-classical receptive-field effects. Nat. Neurosci. 2, 79–87 (1999). 10.1038/4580

2. Friston, K. J. A theory of cortical responses. Philos. Trans. R. Soc. Lond. B Biol. Sci. 360, 815–836 (2005). 10.1098/rstb.2005.1622

3. Friston, K. J. The free-energy principle: A unified brain theory? Nat. Rev. Neurosci. 11, 127–138 (2010). 10.1038/nrn2787

4. Clark, A. Surfing Uncertainty: Prediction, Action, and the Embodied Mind (Oxford Univ. Press, Oxford, 2015). 10.1093/acprof:oso/9780190217013.001.0001

5. Friston, K. J. & Kiebel, S. Predictive coding under the free-energy principle. Philos. Trans. R. Soc. Lond. B Biol. Sci. 364, 1211–1221 (2009). 10.1098/rstb.2008.0300

6. Feldman, H. & Friston, K. J. Attention, uncertainty, and free-energy. Front. Hum. Neurosci. 4, 215 (2010). 10.3389/fnhum.2010.00215

7. Bekinschtein, T. A., Dehaene, S., Rohaut, B., Tadel, F., Cohen, L., & Naccache, L. (2009). Neural signature of the conscious processing of auditory regularities. PNAS 106(5), 1672–1677. 10.1073/pnas.0809667106

8. Wacongne, C. et al. Evidence for a hierarchy of predictions and prediction errors in human cortex. PNAS 108, 20754–20759 (2011). 10.1073/pnas.1117807108

9. Bonetti, L., Fernández-Rubio, G., Carlomagno, F. et al. Spatiotemporal brain hierarchies of auditory memory recognition and predictive coding. Nat Commun 15, 4313 (2024). 10.1038/s41467-024-48302-4

10. Fernández-Rubio, G., Vuust, P., Kringelbach, M.L., et al. The neurophysiology of healthy and pathological aging: a comprehensive systematic review. Brain Struct Funct 230, 146 (2025). 10.1007/s00429-025-03012-5

11. Heng, J. G. et al. Understanding music and aging through the lens of Bayesian inference. Neuroscience & Biobehavioral Reviews 163, 105768 (2024). 10.1016/j.neubiorev.2024.105768

12. Malvaso, C. et al. FREQ-NESS reveals age-related differences in frequency-resolved brain networks during auditory recognition and resting state. bioRxiv (2025) 10.1101/2025.06.30.662094

13. Costa, M., Vuust, P., Kringelbach, M. L. & Bonetti, L. EEG correlates of auditory ShortLTerm memory and dissimilarity perception in young and older adults. European Journal of Neuroscience 61, e70166 (2025). 10.1111/ejn.70166

14. Criscuolo, A., Schwartze, M., Bonetti, L. & Kotz, S. A. Aging impacts basic auditory and timing processes. European Journal of Neuroscience 61, e70031 (2025). 10.1111/ejn.70031

15. Ruzzoli, M., Pirulli, C., Brignani, D., Maioli, C. & Miniussi, C. Sensory memory during physiological aging indexed by mismatch negativity (MMN). Neurobiology of Aging 33, 625.e21–625.e30 (2011). 10.1016/j.neurobiolaging.2011.03.021

16. Cheng, C. H., Hsu, W. & Lin, Y. Y. Effects of physiological aging on mismatch negativity: A meta-analysis. Int. J. Psychophysiol. 90, 165–171 (2013). 10.1016/j.ijpsycho.2013.06.026

17. Cooper, R. J., Todd, J., McGill, K. & Michie, P. T. Auditory sensory memory and the aging brain: A mismatch negativity study. Neurobiol. Aging 27, 752–762 (2006). 10.1016/j.neurobiolaging.2005.03.012

18. Näätänen, R. et al. The mismatch negativity: An index of cognitive decline in neuropsychiatric and neurological diseases and in ageing. Brain 134, 3435–3453 (2011). 10.1093/brain/awr064

19. Chan, J. S. et al. Predictive coding over the lifespan: Increased reliance on perceptual priors in older adults — A magnetoencephalography and dynamic causal modeling study. Front. Aging Neurosci. 13, 631599 (2021). 10.3389/fnagi.2021.631599

20. Wolpe, N., et al. Ageing increases reliance on sensorimotor prediction through structural and functional differences in frontostriatal circuits. Nat Commun 7(1). (2016). 10.1038/ncomms13034

21. Hsu, Y. F., Waszak, F., Strömmer, J. & Hämäläinen, J. A. Human brain ages with hierarchy-selective attenuation of prediction errors. Cereb. Cortex 31, 2156–2168 (2021). 10.1093/cercor/bhaa352

22. Czigler, I., Csibra, G. & Csontos, A. Age and inter-stimulus interval effects on event-related potentials to frequent and infrequent auditory stimuli. Biol. Psychol. 33, 195–206 (1992). 10.1016/0301-0511(92)90031-O

23. Alain, C. & Woods, D. L. Age-related changes in processing auditory stimuli during visual attention: Evidence for deficits in inhibitory control and sensory memory. Psychol. Aging 14, 507–519 (1999). 10.1037/0882-7974.14.3.507

24. Cooper, R. J., Todd, J., McGill, K. & Michie, P. T. Auditory sensory memory and the aging brain: A mismatch negativity study. Neurobiol. Aging 27, 752–762 (2006). 10.1016/j.neurobiolaging.2005.03.012

25. Jääskeläinen, I. P., Varonen, R., Näätänen, R. & Pekkonen, E. Decay of cortical preattentive sound discrimination in middle-age. NeuroReport 10, 123–126 (1999).

26. Gunter, T. C., Jackson, J. L., & Mulder, G. Focussing on aging: an electrophysiological exploration of spatial and attentional processing during reading. Biological Psychology, 43(2), 103–145. (1996). 10.1016/0301-0511(95)05180-5

27. Mueller, V., Brehmer, Y., Von Oertzen, T., Li, S. C., & Lindenberger, U. Electrophysiological correlates of selective attention: A lifespan comparison. BMC Neuroscience, 9(1). (2008). 10.1186/1471-2202-9-18

28. Pekkonen, E., Jousmäki, V., Partanen, J., & Karhu, J. Mismatch negativity area and age-related auditory memory. Electroencephalography and Clinical Neurophysiology, 87(5), 321–325. (1993). 10.1016/0013-4694(93)90185-x

29. Kazmerski, V. A., Friedman, D. & Ritter, W. Mismatch negativity during attend and ignore conditions in Alzheimer’s disease. Biological Psychiatry 42, 382–402 (1997). 10.1016/S0006-3223(96)00344-7

30. Amenedo, E. & Diaz, F. Automatic and effortful processes in auditory memory reflected by event-related potentials. Age-related findings. Electroencephalography and Clinical Neurophysiology 108, 361–369. (1998). 10.1016/s0168-5597(98)00007-0

31. Ruzzoli, M., Pirulli, C., Brignani, D., Maioli, C., Miniussi, C. Sensory memory during physiological aging indexed by mismatch negativity (MMN). Neurobiology of Aging 33 (625), e621–e630. (2012). 10.1016/j.neurobiolaging.2011.03.021

32. Verleger, R., Neukäter, W., Kömpf, D. & Vieregge, P. On the reasons for the delay of P3 latency in healthy elderly subjects. Electroencephalography and Clinical Neurophysiology 79, 488–502 (1991). 10.1016/0013-4694(91)90168-4

33. Polich, J. Theoretical overview of P3a and P3b. In Detection of Change: Event-related Potential and fMRI Findings (Polich, J. ed.) 83–98 (Kluwer Academic, 2003). 10.1007/978-1-4615-0294-4_5

34. Polich, J. Updating P300: An integrative theory of P3a and P3b. Clin. Neurophysiol. 118, 2128–2148 (2007). 10.1016/j.clinph.2007.04.019

35. Kompus, K., Volehaugen, V., Todd, J. & Westerhausen, R. Hierarchical modulation of auditory prediction error signaling is independent of attention. Cogn. Neurosci. 11, 132–142 (2020). 10.1080/17588928.2019.1648404

36. Cohen, M. X. (2014). Analyzing Neural Time Series Data: Theory and Practice. MIT Press.

37. Luck, S. J. (2014). An Introduction to the Event-Related Potential Technique, second edition. MIT Press.

38. Garrido, M. I., Kilner, J. M., Stephan, K. E. & Friston, K. J. The mismatch negativity: A review of underlying mechanisms. Clin. Neurophysiol. 120, 453–463 (2009). 10.1016/j.clinph.2008.11.029

39. Fitzgerald, K. & Todd, J. Making sense of mismatch negativity. Front. Psychiatry 11, 468 (2020). 10.3389/fpsyt.2020.00468

40. Winkler, I. Interpreting the mismatch negativity. J. Psychophysiol. 21, 147–163 (2007). 10.1027/0269-8803.21.34.147

41. Escera, C. & Corral, M. J. Role of mismatch negativity and novelty-P3 in involuntary auditory attention. J. Psychophysiol. 21, 251–264 (2007). 10.1027/0269-8803.21.34.251

42. Justo-Guillén, E. et al. (2019). Auditory mismatch detection, distraction, and attentional reorientation (MMN-P3a-RON) in neurological and psychiatric disorders: A review. International Journal of Psychophysiology 146, 85–100. 10.1016/j.ijpsycho.2019.09.010

43. Munka, L., & Berti, S. (2006). Examining task-dependencies of different attentional processes as reflected in the P3a and reorienting negativity components of the human event-related brain potential. Neuroscience Letters, 396(3), 177–181. 10.1016/j.neulet.2005.11.035

44. Chennu, S. et al. Expectation and attention in hierarchical auditory prediction. J. Neurosci. 33, 11194–11205 (2013). 10.1523/JNEUROSCI.0114-13.2013

45. Baars, B. J. Global workspace theory of consciousness: Toward a cognitive neuroscience of human experience. Prog. Brain Res. 150, 45–53 (2005). 10.1016/S0079-6123(05)50004-9

46. Dehaene, S. Towards a cognitive neuroscience of consciousness: Basic evidence and a workspace framework. Cognition 79, 1–37 (2001). 10.1016/S0010-0277(00)00123-2

47. Whyte, C. J. Integrating the global neuronal workspace into the framework of predictive processing: Towards a working hypothesis. Conscious. Cogn. 73, 102763 (2019). 10.1016/j.concog.2019.102763

48. Bonetti, L. et al. BROADLNESS uncovers DualLStream mechanisms underlying predictive coding in auditory memory networks. Advanced Science 12, e07878 (2025). 10.1002/advs.202507878

49. Bonetti, L. et al. Shared and modality-specific brain networks underlying predictive coding of temporal sequences. bioRxiv (Cold Spring Harbor Laboratory) (2025) 10.1101/2025.07.16.665207

50. Del Giudice, M. Effective Dimensionality: a tutorial. Multivariate Behavioral Research 56, 527–542 (2020). 10.1080/00273171.2020.1743631

51. Grady, C. L. (2012). The cognitive neuroscience of ageing. Nature Reviews Neuroscience, 13(7), 491–505. 10.1038/nrn3256

52. Quiroga-Martinez, D. R. et al. Correction: Decoding reveals the neural representation of perceived and imagined musical sounds. PLoS Biology 23, e3003445 (2025). 10.1371/journal.pbio.3003445

53. Bonetti, L. et al. Rapid encoding of musical tones discovered in whole-brain connectivity. NeuroImage 245, 118735 (2021). 10.1016/j.neuroimage.2021.118735

54. Bonetti, L. et al. Spatiotemporal whole-brain activity and functional connectivity of melodies recognition. Cerebral Cortex 34, (2024). 10.1093/cercor/bhae320

55. Bonetti, L. et al. Brain recognition of previously learned versus novel temporal sequences: a differential simultaneous processing. Cerebral Cortex 33, 5524–5537 (2022). 10.1093/cercor/bhac439

56. Fernández-Rubio, G., Brattico, E., Kotz, S.A. et al. Magnetoencephalography recordings reveal the spatiotemporal dynamics of recognition memory for complex versus simple auditory sequences. Commun Biol 5, 1272 (2022). 10.1038/s42003-022-04217-8

57. Fernández-Rubio, G., Carlomagno, F., Vuust, P., Kringelbach, M. L. & Bonetti, L. Associations between abstract working memory abilities and brain activity underlying long-term recognition of auditory sequences. PNAS Nexus 1, pgac216 (2022). 10.1093/pnasnexus/pgac216

58. Hsu, Y.-F., Hämäläinen, J. A., Waszak, F., & Vissers, M. Longitudinal evidence for attenuated local–global deviance detection as a precursor of working memory decline. eNeuro, 10(8). (2023). 10.1523/ENEURO.0156-23.2023

59. Donchin, E. & Coles, M. G. H. Is the P300 component a manifestation of context updating? Behav. Brain Sci. 11, 357–374 (1988). 10.1017/S0140525X00058027

60. Bonetti, L. et al. Age-related neural changes underlying long-term recognition of musical sequences. Communications Biology 7, 1036 (2024). 10.1038/s42003-024-06587-7

61. Fernández-Rubio, G. et al. Investigating the impact of age on auditory short-term, long-term, and working memory. Psychology of Music 52, 187–198 (2023). 10.1177/0305735623118340

62. Bonetti, L. et al. Working memory predicts LongLTerm recognition of auditory sequences: dissociation between confirmed predictions and prediction errors. Scandinavian Journal of Psychology 66, 842–853 (2025). 10.1111/sjop.13124

63. Friedman, D., Kazmerski, V. A., & Cycowicz, Y. M. (1998). Effects of aging on the novelty P3 during attend and ignore oddball tasks. Psychophysiology, 35(5), 508–520. 10.1017/S0048577298970664

64. Gaeta, H., Friedman, D., Ritter, W., & Cheng, J. (1998). An event-related potential study of age-related changes in sensitivity to stimulus deviance. Neurobiology of Aging, 19(5), 447–459. 10.1016/S0197-4580(98)00087-6

65. Cabeza, R. Hemispheric asymmetry reduction in older adults: The HAROLD model. Psychology and Aging 17, 85–100 (2002). 10.1037//0882-7974.17.1.85

66. May, P. J. C. & Tiitinen, H. Mismatch negativity (MMN), the deviance-elicited auditory deflection, explained. Psychophysiology 47, 66–122 (2010). 10.1111/j.1469-8986.2009.00856.x

67. Baldeweg, T. ERP repetition effects and mismatch negativity generation. J. Psychophysiol. 21, 204–213 (2007). 10.1027/0269-8803.21.34.204

68. Herrmann, B., Henry, M. J., Johnsrude, I. S. & Obleser, J. Altered temporal dynamics of neural adaptation in the aging human auditory cortex. Neurobiol. Aging 45, 10–22 (2016). 10.1016/j.neurobiolaging.2016.05.006

69. Willmore, B. D. B. & King, A. J. Adaptation in auditory processing. Physiol. Rev. 103, 1025–1058 (2023). 10.1152/physrev.00011.2022

70. Sur, S. & Golob, E. J. Neural correlates of auditory sensory memory dynamics in the aging brain. Neurobiol. Aging 88, 128–136 (2020). 10.1016/j.neurobiolaging.2019.12.020

71. Rutiku, R., Fiscone, C., Massimini, M. & Sarasso, S. Assessing mismatch negativity (MMN) and P3b within-individual sensitivity — A comparison between the local–global paradigm and two specialized oddball sequences. Eur. J. Neurosci. 59, 842–859 (2024). 10.1111/ejn.16302

72. Dehaene, S. Towards a cognitive neuroscience of consciousness: Basic evidence and a workspace framework. Cognition 79, 1–37 (2001). 10.1016/S0010-0277(00)00123-2

73. King, J.-R. et al. (2014). Single-trial decoding of auditory novelty responses facilitates the detection of residual consciousness. NeuroImage 83, 726–738. 10.1016/j.neuroimage.2013.07.013

74. Marti, S., King, J.-R., & Dehaene, S. (2014). Time-resolved decoding of two processing chains during dual-task interference. PLoS ONE 9(2), e89386. 10.1371/journal.pone.0089386

75. Niedernhuber, M., Candia-Rivera, D., Pokorny, C., Chennu, S., & Bekinschtein, T. A. (2022). Sensory target detection at local and global timescales: A novel dissociation between conscious access and task performance. Journal of Neuroscience 42(50), 9424–9437. 10.1523/JNEUROSCI.0233-22.2022

76. Candia-Rivera, D. et al. Conscious processing of global and local auditory irregularities causes differentiated heartbeat-evoked responses. eLife 12, (2023). 10.7554/eLife.75352

77. Bastos, A. M. et al. Canonical microcircuits for predictive coding. Neuron 76, 695–711 (2012). 10.1016/j.neuron.2012.10.038

78. Peirce, J. W., Gray, J. R., Simpson, S., MacAskill, M. R., Höchenberger, R., Sogo, H., Kastman, E., Lindeløv, J. (2019). PsychoPy2: experiments in behavior made easy. Behavior Research Methods. 10.3758/s13428-018-01193-y

79. Wasano, K., Kaga, K., & Ogawa, K. (2021). Patterns of hearing changes in women and men from denarians to nonagenarians. The Lancet Regional Health – Western Pacific, 9, 100131. 10.1016/j.lanwpc.2021.100131

80. Taulu, S. & Simola, J. Spatiotemporal signal space separation method for rejecting nearby interference in MEG measurements. Phys. Med. Biol. 51, 1759–1768 (2006). 10.1088/0031-9155/51/7/008

81. Woolrich, M., Hunt, L., Groves, A. & Barnes, G. MEG beamforming using Bayesian PCA for adaptive data covariance matrix regularization. Neuroimage 57, 1466–1479 (2011). 10.1016/j.neuroimage.2011.04.041

82. Oostenveld, R., Fries, P., Maris, E. & Schoffelen, J.-M. FieldTrip: open source software for advanced analysis of MEG, EEG, and invasive electrophysiological data. Comput. Intell. Neurosci. 2011, 1–9 (2011). 10.1155/2011/156869

83. Woolrich, M. W. et al. Bayesian analysis of neuroimaging data in FSL. Neuroimage 45, 173–186 (2009). 10.1016/j.neuroimage.2008.10.055

84. Penny, W. D., Friston, K. J., Ashburner, J. T., Kiebel, S. J. & Nichols, T. E. *Statistical Parametric Mapping: The analysis of Functional Brain Images*. (Elsevier, London, 2011). 10.1016/B978-0-12-372560-8.X5000-1

85. Mantini, D. et al. A signal-processing pipeline for magnetoencephalography resting-state networks. Brain Connect. 1, 49–59 (2011). 10.1089/brain.2011.0001

86. Wang, G., Teng, C., Li, K., Zhang, Z. & Yan, X. The removal of EOG artifacts from EEG signals using independent component analysis and multivariate empirical mode decomposition. IEEE J. Biomed. Health Inform. 20, 1301–1308 (2015). 10.1109/JBHI.2015.2450196

87. Brookes, M. J. et al. Beamformer reconstruction of correlated sources using a modified source model. Neuroimage 34, 1454–1465 (2007). 10.1016/j.neuroimage.2006.11.012

88. Huang, M., Mosher, J. C. & Leahy, R. A sensor-weighted overlapping-sphere head model and exhaustive head model comparison for *MEG*. Phys. Med. Biol. 44, 423 (1999). 10.1088/0031-9155/44/2/010

89. Hillebrand, A. & Barnes, G. R. Beamformer analysis of MEG data. Int. Rev. Neurobiol. 68, 149–171 (2005). 10.1016/S0074-7742(05)68006-3

90. Nolte, G. The magnetic lead field theorem in the quasi-static approximation and its use for magnetoencephalography forward calculation in realistic volume conductors. Phys. Med. Biol. 48, 3637–3652 (2003). 10.1088/0031-9155/48/22/002

91. Huang, M., Mosher, J. C. & Leahy, R. A sensor-weighted overlapping-sphere head model and exhaustive head model comparison for *MEG*. Phys. Med. Biol. 44, 423 (1999). 10.1088/0031-9155/44/2/010

92. Gross, J. et al. Good practice for conducting and reporting MEG research. Neuroimage 65, 349–363 (2013). 10.1016/j.neuroimage.2012.10.001

93. Afnan, J. et al. Validating MEG source imaging of resting state oscillatory patterns with an intracranial EEG atlas. Neuroimage 274, 120158 (2023). 10.1016/j.neuroimage.2023.120158

94. Sauseng, P., Griesmayr, B., Freunberger, R. & Klimesch, W. Control mechanisms in working memory: a possible function of EEG theta oscillations. Neurosci. Biobehav Rev. 34, 1015–1022 (2010). 10.1016/j.neubiorev.2009.12.006

95. Hall, E. L., Woolrich, M.W., Thomaz, C.E., Morris, P.G. & Brookes, M. J. Using variance information in magnetoencephalography measures of functional connectivity. Neuroimage 67, 203–212 (2013). 10.1016/j.neuroimage.2012.11.011

96. Luckhoo, H. T., Brookes, M. J. & Woolrich, M. W. Multi-session statistics on beamformed MEG data. Neuroimage 95, 330–335 (2014). 10.1016/j.neuroimage.2013.12.026

97. Van Veen, B. D., van Drongelen, W., Yuchtman, M. & Suzuki, A. Localization of brain electrical activity via linearly constrained minimum variance spatial filtering. IEEE Trans. Biomed. Eng. 44, 867–880 (1997). 10.1109/10.623056

98. Haufe, S. et al. On the interpretation of weight vectors of linear models in multivariate neuroimaging. Neuroimage 87, 96–110 (2013). 10.1016/j.neuroimage.2013.10.067

99. Wall, M. E., Rechtsteiner, A. & Rocha, L. M. in Kluwer Academic Publishers eBooks, 91–109 (2005). 10.1007/0-306-47815-3_5

100. Maris, E. & Oostenveld, R. Nonparametric statistical testing of EEG– and MEG-data. J. Neurosci. Methods 164, 177–190 (2007). 10.1016/j.jneumeth.2007.03.024

101. Marwan, N., Wessel, N., Meyerfeldt, U., Schirdewan, A. & Kurths, J. Recurrence-plot-based measures of complexity and their application to heart-rate-variability data. Phys Rev E Stat Phys Plasmas Fluids Relat Interdiscip Topics 66, (2002). 10.1103/PhysRevE.66.026702

102. Marwan, N., Carmenromano, M., Thiel, M. & Kurths, J. Recurrence plots for the analysis of complex systems. Phys. Rep. 438, 237–329 (2007). 10.1016/j.physrep.2006.11.001

103. Benjamini, Y., & Hochberg, Y. (1995). Controlling the False Discovery Rate: A Practical and Powerful Approach to Multiple Testing. J. R. Stat. Soc. Ser. B Stat. Methodol., 57(1), 289–300. 10.1111/j.2517-6161.1995.tb02031.x

104. Sinaga, K. P. & Yang, M.-S. Unsupervised K-Means clustering algorithm. IEEE Access 8, 80716–80727 (2020). 10.1109/ACCESS.2020.2988796

105. Garcia-Dias R, Vieira S, Lopez Pinaya W. H., Mechelli A. Clustering analysis. In: A. Mechelli & S. Vieira. (Eds.). Machine learning: Methods and applications to brain disorders. Cambridge, USA: Academic Press (2019). 10.1016/B978-0-12-815739-8.00013-4

106. Zoubi, M. B. A.- & Rawi, M. A. An efficient approach for computing silhouette coefficients. J. Comput. Sci. 4, 252–255 (2008). 10.3844/jcssp.2008.252.255

107. Bonetti, L. et al. Hierarchical predictive processing in younger and older adults – MEG dataset [Dataset]. Zenodo. (2026). 10.5281/zenodo.18231641

