## Supplementary materials for "Network reconfiguration preserves prediction error signalling in the aging brain"

*Mathias Houe Andersen _1, 2, 3_*_,_ Gemma Fernández-Rubio _1,_ David R. Quiroga-Martinez _1,4,_*

*Mattia Rosso _1,5,_ Mathias Klarlund _1,_* *Kit Melissa Larsen _2,_ Hartwig Roman Siebner _2,3,6,_*

*Morten L. Kringelbach _1,7,8,_ Peter Vuust _1,_ Leonardo Bonetti _1,7,8_**

^1^ Center for Music in the Brain, Department of Clinical Medicine, Aarhus University & The Royal Academy of Music, Aarhus/Aalborg, Denmark

^2^ Danish Research Centre for Magnetic Resonance, Department of Radiology and Nuclear Medicine, Copenhagen University Hospital – Amager and Hvidovre, Hvidovre, Denmark

^3^ Faculty of Health and Medical Sciences, University of Copenhagen, Copenhagen, Denmark

^4^ Department of Psychology, University of Copenhagen, Copenhagen, Denmark

^5^ IPEM Institute for Systematic Musicology, Ghent University, Ghent, Belgium

^6^ Department of Neurology, Copenhagen University Hospital Bispebjerg and Frederiksberg, Copenhagen, Denmark

^7^ Department of Psychiatry, University of Oxford, Oxford, United Kingdom

^8^ Centre for Eudaimonia and Human Flourishing, Linacre College, University of Oxford, Oxford, United Kingdom

ORCID:

https://orcid.org/0009-0003-2513-5251

https://orcid.org/0000-0001-9983-3819

**Supplementary figures**

The same supplementary figures are available in high resolution at the following link:

https://doi.org/10.5281/zenodo.18508401

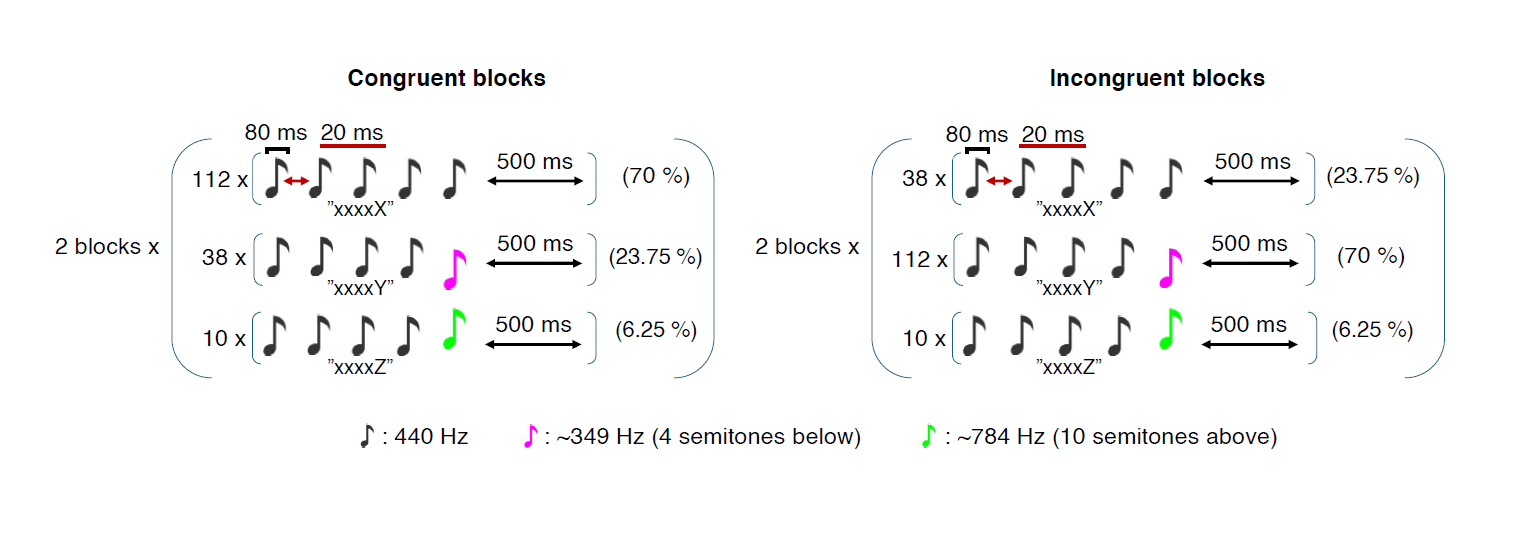

**Fig. S1 | The auditory local-global paradigm.**

Each trial consisted of patterns of five tones. The duration of each tone was 80 ms separated by 20 ms of silence yielding a stimulus onset asynchrony of 100 ms. Each five-tone pattern was followed by a 500 ms inter-trial interval. Tone patterns could be either five identical tones (local standards) or sequences in which the fifth tone differed in pitch (local deviants). Complementing this experimental fifth tone that induces local sensory violations, a target tone requiring a button press was also rarely presented to ensure continued attention towards the paradigm. Blocks manipulated prediction violations at two hierarchical levels. In congruent blocks, local standards occurred most frequently (70%), local experimental deviants occurred less often (23.75%), and target tones appeared rarely (6.25%). In incongruent blocks, the probabilities of local standards and local deviants were switched, creating a global violation in pattern frequency, while target probability remained constant. Participants completed two congruent and two incongruent blocks in randomized order. Three pitch levels were used (standard tones: 440 Hz; deviating tones: 4 semitones below standards, ~349 Hz; target tones: 10 semitones above standards, ~784 Hz). This design elicits local prediction errors at the tone level and global prediction errors at the pattern level, while maintaining attention through infrequent targets.

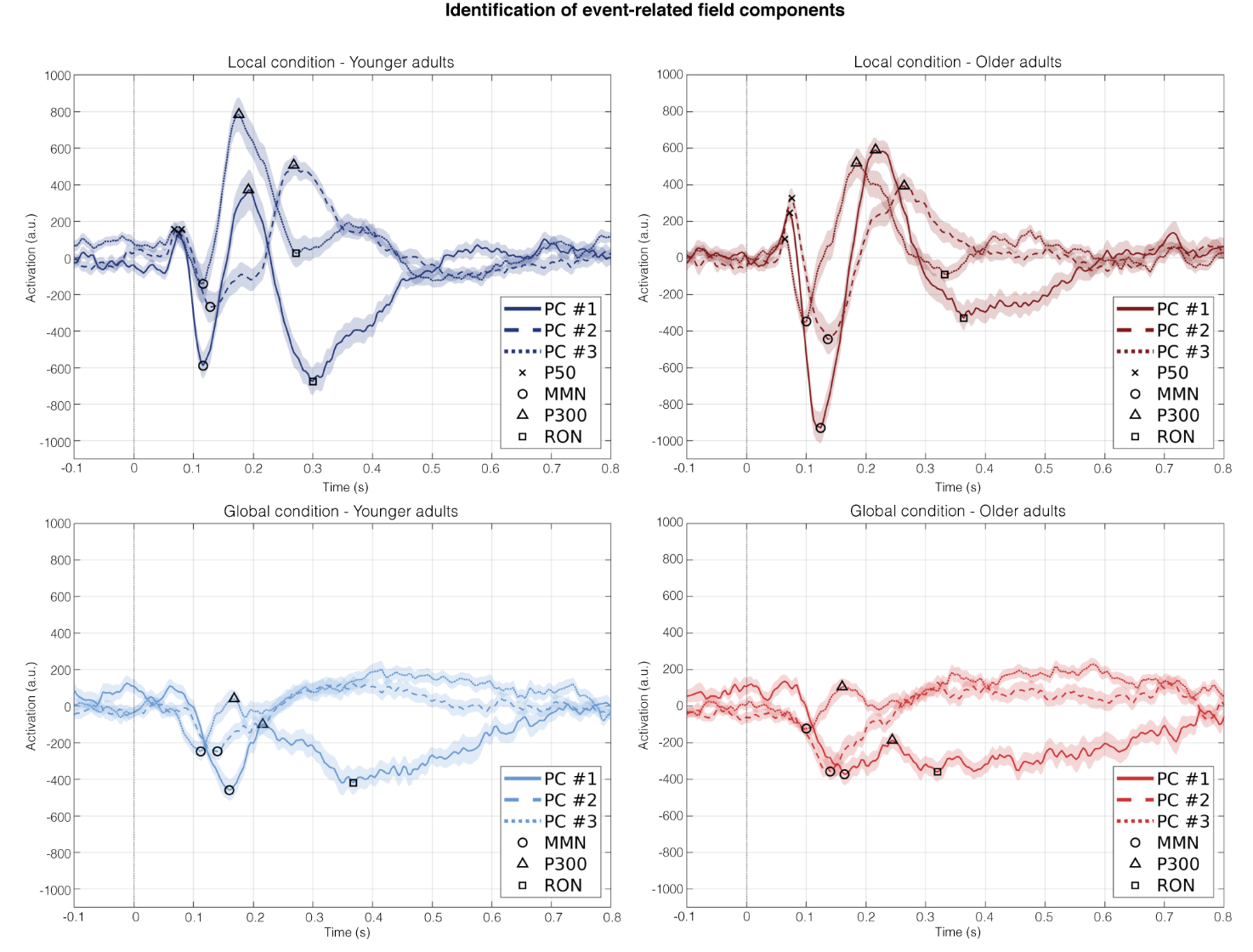

**Figure S2 | Event-related field components identification.**

This figure illustrates the identification of canonical event-related field (ERF) components from group-mean brain network time series derived from principal component (PC) decomposition. Mean activation time courses (μ ± SE) are shown separately for younger and older participants under local and global conditions, with PC #1–3 distinguished by line style. ERF components were detected by extracting the most positive or negative peak within predefined latency windows: for the local condition, P50 (50 – 100 ms, positive), MMN (100 – 200 ms, negative), P300 (150 – 300 ms, positive), and RON (250 – 400 ms, negative); for the global condition, MMN (100 – 200 ms, negative), P300 (150 – 250 ms, positive), and RON (250 – 400 ms, negative). Peak latencies are marked with symbols on the corresponding time series. Components that were not reliably identifiable were excluded a priori (e.g., RON in PC #2 for the local condition, P300 in PC #2 for the global condition, and P50 in the global condition), and global RON was only quantified for PC #1. These peak estimates form the basis for the latency, amplitude, and temporal order summary reported in Table S4.

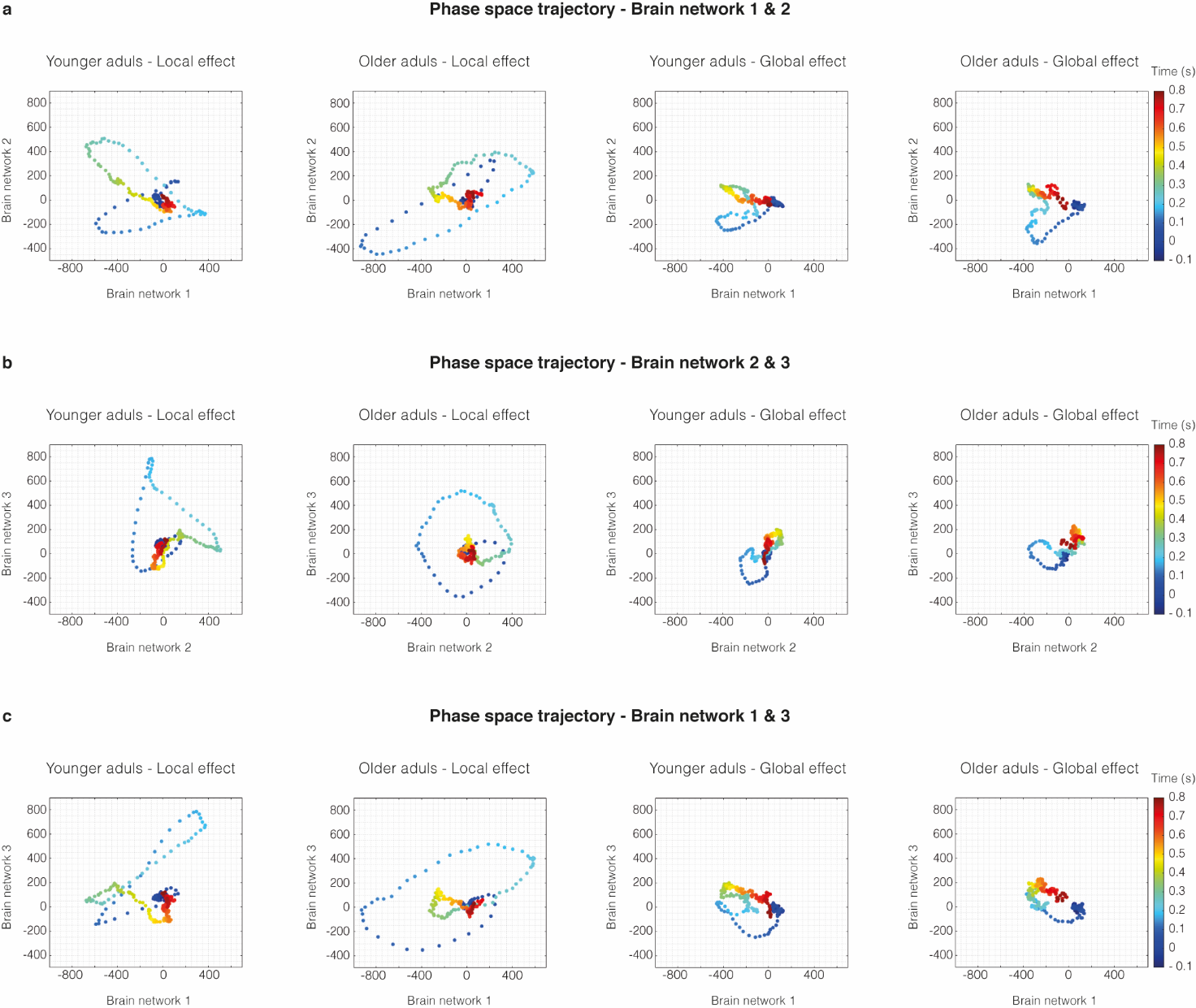

**Figure S3 | Phase space trajectories in two dimensions.**

Phase space trajectories averaged across participants for each experimental condition and age group, obtained by embedding the time series of the three primary BROAD-NESS networks into a two-dimensional state space in all three combinations possible. Each dot represents the state of the brain at a given time point, colour-coded by time (in seconds). Across the 3 different comparisons (PC #1 vs. PC #2, PC #2 vs. PC #3 and PC #1 vs. PC #3), in the local condition younger adults show a more sharply defined time specific coactivation and deactivation of singular networks compared to slower and less distinct transitions observed among older adults. Both groups display a more concentrated time specific activation and deactivation of individual networks in the local compared to the global condition.

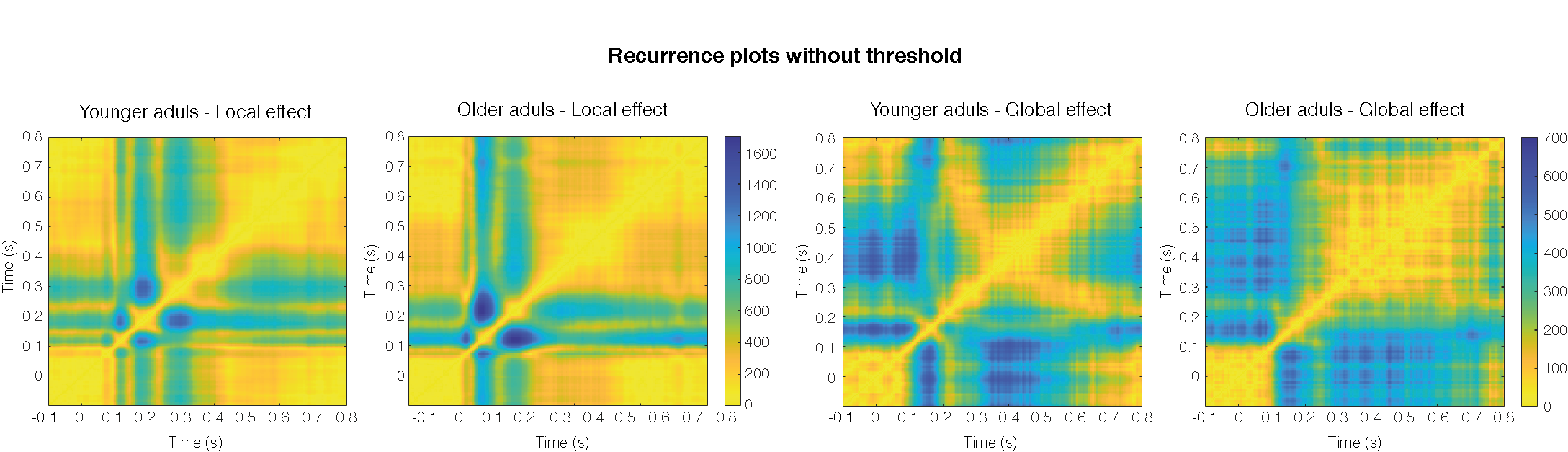

**Figure S4 | Recurrence plots without threshold.**

Recurrence plots computed from the three-dimensional phase space trajectories in Fig. 3b. The matrices are averaged across participants, depicting pairwise Euclidean distances between time points. Blue indicates larger distances (i.e., less similar brain states), while yellow indicate greater similarity and temporal recurrence. Across both conditions, younger adults show less recurrence of the three brain networks across time compared to older adults indicating more exploratory and less static temporal dynamics regarding the reutilisation of brain networks.

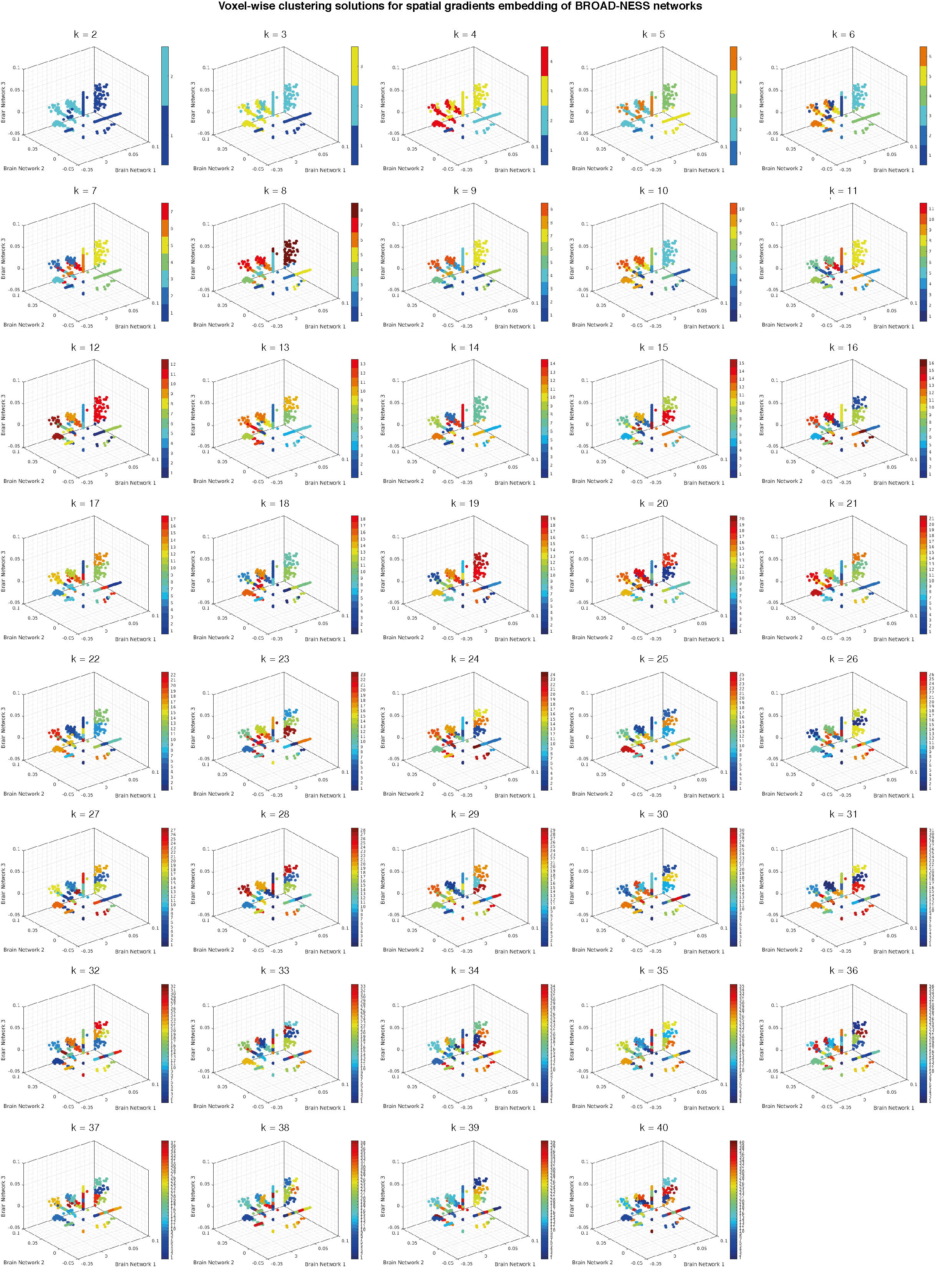

**Figure S5 | Voxel-wise clustering solutions for spatial gradients embedding of BROAD-NESS-derived brain networks (k = 2 to 40).**

Each subplot shows a 3D embedding of brain voxels based on their contribution to the three principal BROAD-NESS networks, with colour-coded clusters derived from unsupervised k-means clustering for values of k ranging from 1 to 40. The embedding reflects how individual voxels co-participate across networks, revealing patterns of overlap, selectivity, or opposition. The full panel allows visual comparison across clustering solutions to identify consistent subdivisions the brain voxels across multiple runs of clustering analysis. The optimal number of clusters, determined through silhouette scores and repetition stability (corresponding to k = 16, is reported both here and in higher detail in the main text.

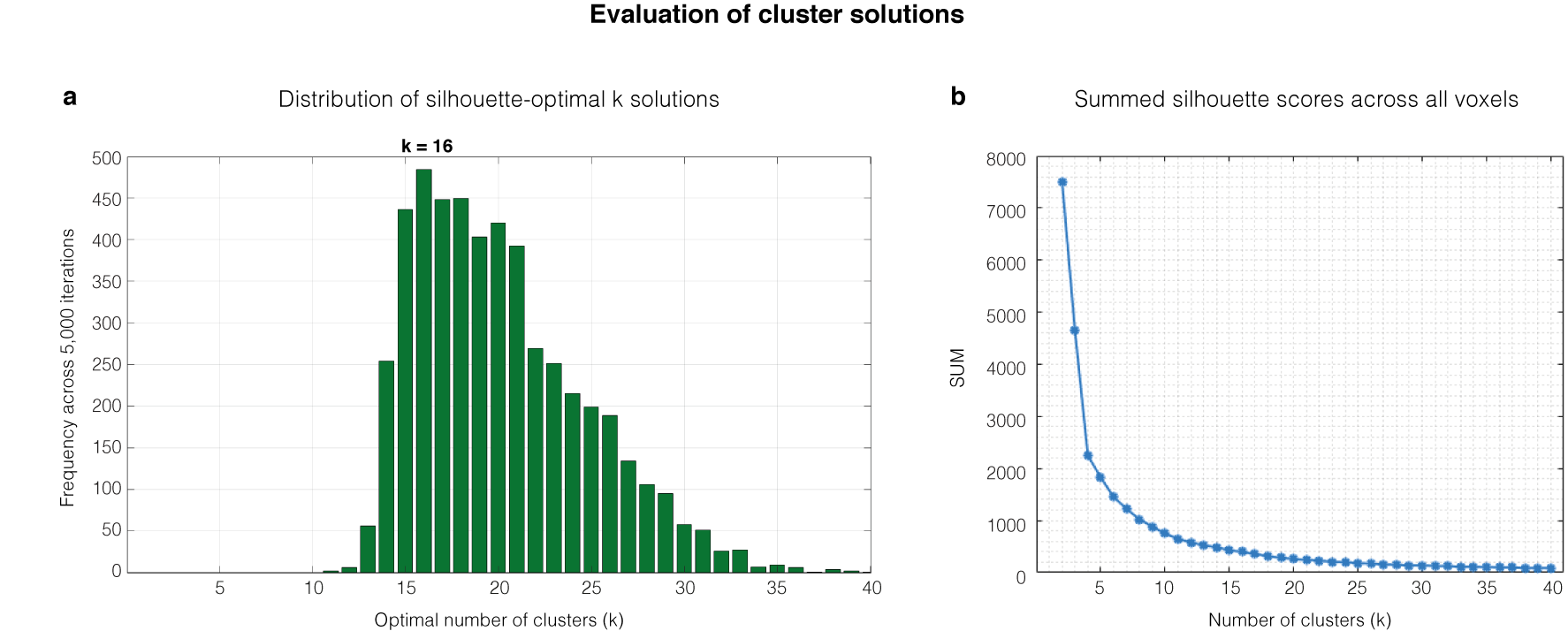

**Figure S6 | Evaluation of cluster solutions.**

**a**, Histogram of the frequency with which each *k* was selected as silhouette-optimal across 5,000 repetitions for each tested cluster solution (*k* = 2 to 40) of the spatial gradients embedding of BROAD-NESS-derived brain networks. The peak at *k* = 16 confirms the optimal number of spatial clusters, based on maximum silhouette coefficient consistency. **b**, The plot shows the within-cluster sum of distances (SUM; k-means objective) as a function of the number of clusters *k* (here, *k* = 2 to 40), computed from voxel embeddings in principal-component weight space after thresholding and z-scoring. For each *k*, k-means clustering was run with 100 replicates with different random initializations, and the best solution with the lowest within-cluster dispersion across replicates was retained. SUM decreases monotonically as *k* increases since additional clusters allow a progressively finer partitioning of the embedding space.

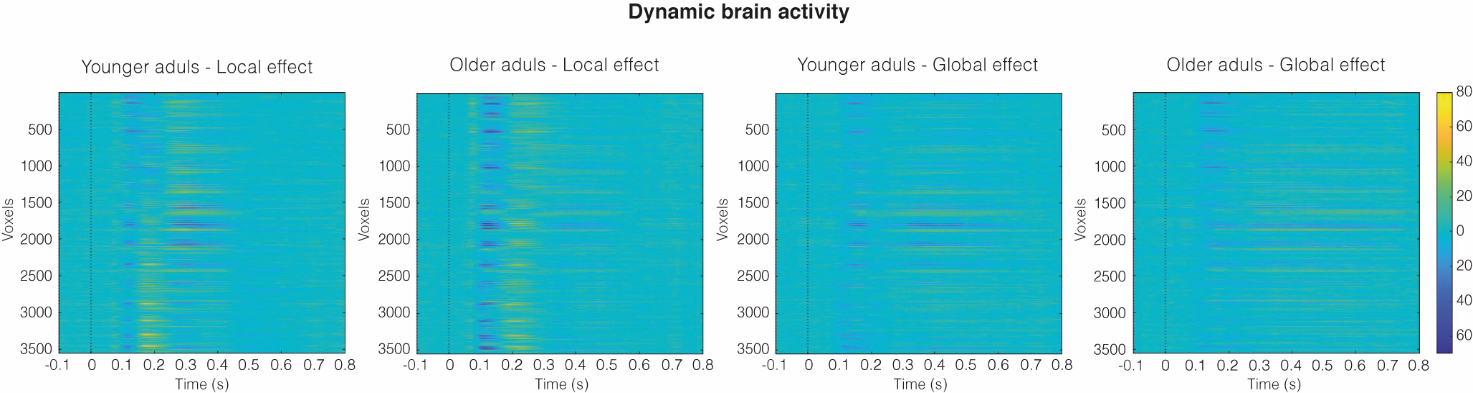

**Figure S7 | Dynamic brain activity.**

The figure shows voxel-wise dynamic brain activity derived directly from the pre-processed data (voxels × time × condition × participants). Data was averaged across each group in each condition. Each row in the heat map corresponds to one reconstructed brain voxel in the 3,559 voxel grid used, and each column corresponds to a time point covering 4 ms due to the 250 Hz resampling rate. The figure highlights condition differences in the temporal activation of brain voxels, where voxels in the local condition are activated in a more concentrated and time specific manner compared to the less punctual activation of voxels in the global condition. Furthermore, older adults display a stronger deactivation and activation of voxels in the 100 – 250 ms time range compared to younger adults. No statistical testing procedures were performed. The figure simply acts as an initial inspection of the voxel activations across time before deriving brain networks from the BROAD-NESS methodology.

**Supplementary Tables**

The supplementary tables are available at the following link:

https://www.doi.org/10.5281/zenodo.18390836

**Table S1 | MNI coordinates of brain voxels from the three main brain networks.**

This table reports the results of the voxels corresponding to the three main brain networks derived from BROAD-NESS in the main analysis. For each voxel, the table provides its progressive index and MNI coordinates (X, Y, Z), along with its corresponding values (spatial activation pattern) on PC #1, PC #2 and PC #3.

**Table S2 | Labels of brain areas within the three main brain networks.**

This table summarizes voxel-level contributions from each brain network after mapping MNI-space coordinates to anatomical regions defined by the AAL 3.1 atlas^[1]^. Voxels were first separated into positive and negative contributions based on the sign of the network loading. Each voxel was assigned an atlas label using exact voxel overlap when possible; voxels falling in background or unlabelled regions were iteratively associated with the nearest non-zero atlas label within a predefined voxel radius. Counts are reported per brain area and stratified by the radius at which a valid label was obtained (exact match, 1-voxel shell, 2-voxel shell, 3-voxel shell), with unresolved voxels tracked separately. Distances are defined in atlas voxel space using a Chebyshev neighbourhood metric, where distance is determined by the largest difference in coordinates along any axis. Additional summary tables report the overall distribution of brain area labelling across voxels.

**Table S3 | Statistical analysis of the timeseries from the three main brain networks.**

This table presents the results of statistical analyses conducted independently for the three brain networks that explained the highest variance. Time series of these principal components were analysed in a mixed design with the between-subject factor Group (younger vs. older adults) and the within-subject factor Condition (local vs. global) in the 0 – 800 ms post-stimulus interval. Specifically, we derived three contrast time series per network: (i) a Group main effect (younger minus older, averaging across local and global), (ii) a Condition main effect (global minus local, pooling across age groups), and (iii) a Group × Condition interaction (the difference of the global–local contrast between younger and older adults). For each contrast and each time point, we applied a two-sided cluster-based permutation test over time spanning 5,000 permutations.

**Table S4 | Latency, order and amplitude of event-related field components.**

This table summarizes the peak latency, peak amplitude, and temporal order of event-related field components derived from group-mean time series, reported separately for younger and older participants and for the local and global condition. Event-related field components were quantified from the mean signal (μ) of each principal component (PC), averaged across participants. Component amplitudes were defined within predefined latency windows as either the most positive or negative value depending on the component polarity. For the local condition, P50 (50–100 ms, positive), MMN (100–200 ms, negative), P300 (150–300 ms, positive), and RON (250–400 ms, negative) were analysed. For the global condition, MMN (100–200 ms, negative) and P300 (150–250 ms, positive) were considered. Some event-related field components were not easily identified in the time series and were therefore excluded a priori. This included the RON response for PC #2 in the local condition, and P300 for PC #2 in the global condition. Similarly, in the global condition P50 was not clearly identified in any PC, while the RON response was only identified in PC #1.

**Table S5 | Average 3D phase space trajectory values.**

This table contains time-resolved, group-averaged phase-space coordinates derived from recurrence quantification analysis (RQA) corresponding to both age groups (young, older) and conditions (local, global). Rows represent the time vector followed by the first three principal component (PC) coordinates defining the reconstructed phase space. Columns correspond to successive time points. Values are provided in arbitrary units reflecting PCA-derived component scores. The data enable exact numerical inspection of the reported phase-space plots and allow independent analysis of trajectory geometry.

**Table S6 | MNI coordinates of brain voxels assigned to each cluster in the spatial gradient embedding analysis.**

This table reports the results of the voxel-wise clustering analysis based on the spatial gradient embedding of the three principal BROAD-NESS networks. Voxels were clustered according to their contributions to Network #1, Network #2 and Network #3 (represented by PCA components PC #1, PC # and PC #3, respectively), using an unsupervised k-means clustering approach. The optimal solution identified 16 clusters, each shown in a separate sheet. For each voxel, the table provides its progressive index and MNI coordinates (X, Y, Z), along with its corresponding values (spatial activation pattern) on PC #1, PC #2 and PC #3.

**Table S7 | Labels of brain areas for each cluster in the spatial gradient embedding analysis.**

This table summarizes voxel-level contributions from each brain network after mapping MNI-space coordinates to anatomical regions defined by the AAL 3.1 atlas^[2]^. Each voxel was assigned an atlas label using exact voxel overlap when possible; voxels falling in background or unlabelled regions were iteratively associated with the nearest non-zero atlas label within a predefined voxel radius. Counts are reported per brain area and stratified by the radius at which a valid label was obtained (exact match, 1-voxel shell, 2-voxel shell, 3-voxel shell), with unresolved voxels tracked separately. Distances are defined in atlas voxel space using a Chebyshev neighbourhood metric, where distance is determined by the largest difference in coordinates along any axis. Additional summary tables report the overall distribution of brain area labelling across voxels.

**Table S8 | Minimum and maximum values of brain network loadings in each cluster in the spatial gradient embedding analysis.**

This table reports the results of the voxel-wise clustering analysis based on the spatial gradient embedding of the three principal BROAD-NESS networks. Voxels were clustered according to their contributions to Brain Network #1, Brain Network #2 and Brain Network #3 (represented by PCA components PC #1, PC # and PC #3, respectively), using an unsupervised k-means clustering approach. The optimal solution identified 16 clusters. The table shows the minimum and maximum values of the loadings on each network within each of the 16 clusters. Xmin and Xmax refer to PC #1 loading boundaries, Ymin and Ymax refer to PC #2 loading boundaries, while Zmin and Zmax refer to PC #3 loading boundaries.

**Table S9 | MNI coordinates of brain voxels from the three main brain networks when BROAD-NESS is applied to group and condition separated data.**

This table reports the results of the voxels corresponding to the three main brain networks when BROAD-NESS is applied to group and condition separated data. For each voxel, the table provides its progressive index and MNI coordinates (X, Y, Z), along with its corresponding values (spatial activation pattern) on PC #1, PC #2 and PC #3.

**Table S10 | Labels of brain areas from brain networks when BROAD-NESS is applied to group and condition separated data.**

These four tables summarize voxel-level contributions from each brain network after mapping MNI-space coordinates to anatomical regions defined by the AAL 3.1 atlas. There is one table for each group and condition separated dataset. Voxels were first separated into positive and negative contributions based on the sign of the network loading. Each voxel was assigned an atlas label using exact voxel overlap when possible; voxels falling in background or unlabelled regions were iteratively associated with the nearest non-zero atlas label within a predefined voxel radius. Counts are reported per brain area and stratified by the radius at which a valid label was obtained (exact match, 1-voxel shell, 2-voxel shell, 3-voxel shell), with unresolved voxels tracked separately. Distances are defined in atlas voxel space using a Chebyshev neighbourhood metric, where distance is determined by the largest difference in coordinates along any axis. Additional summary tables report the overall distribution of brain area labelling across voxels.

**Table S11 | Statistical analysis of the timeseries from brain networks when BROAD-NESS is applied to group and condition separated data.**

This table presents the results of statistical analyses conducted independently for the three brain networks within each group-condition-combination when BROAD-NESS was applied to group and condition segregated data. Time series of these principal components were analysed in a mixed design with the between-subject factor Group (younger vs. older adults) and the within-subject factor Condition (local vs. global) in the 0 – 800 ms post-stimulus interval. Specifically, we derived three contrast time series per network: (i) a Group main effect (younger minus older, averaging across local and global), (ii) a Condition main effect (global minus local, pooling across age groups), and (iii) a Group × Condition interaction (the difference of the global–local contrast between younger and older adults). For each contrast and each time point, we applied a two-sided cluster-based permutation test over time spanning 5,000 permutations.

**References**

1. Rolls, E. T., Huang, C. C., Lin, C. P., Feng, J. & Joliot, M. Automated anatomical labelling atlas 3. *NeuroImage* **206**, 116189 (2020). <https://doi.org/10.1016/j.neuroimage.2019.116189>
